# Comparison between slow, anisotropic LE4PD fluctuations and the Principal Component Analysis modes of Ubiquitin

**DOI:** 10.1101/2021.01.06.425617

**Authors:** E. R. Beyerle, M. G. Guenza

## Abstract

Proteins’ biological function and folding mechanisms are often guided by large-scale, slow motions, which involve crossing high energy barriers. In a simulation trajectory, these slow fluctuations are commonly identified using a principal component analysis (PCA). Despite the popularity of this method, a complete analysis of its predictions based on the physics of protein motion has been so far limited. This study formally connects the PCA to a Langevin model of protein dynamics and analyzes the contributions of energy barriers and hydrodynamic interactions to the slow PCA modes of motion. To do so, we introduce an anisotropic extension of the Langevin Equation for Protein Dynamics, called the LE4PD-XYZ, which formally connects to the PCA ‘essential dynamics’. The LE4PD-XYZ is an accurate coarse-grained diffusive method to model protein motion, which describes anisotropic fluctuations in the protein’s alpha-carbons. The LE4PD accounts for hydrodynamic effects and mode-dependent free-energy barriers. This study compares large-scale anisotropic fluctuations identified by the LE4PD-XYZ to the mode-dependent PCA’s predictions, starting from a microsecond-long alpha-carbon molecular dynamics atomistic trajectory of the protein ubiquitin. We observe that the inclusion of free-energy barriers and hydrodynamic interactions has important effects on the identification and timescales of ubiquitin’s slow modes.

## I. INTRODUCTION

Proteins are semi-flexible objects whose function is determined by the combined effect of their three-dimensional structure and local fluctuations^1^. Large-scale, slow motions relevant to the protein’s biological function typically involve crossing high energy barriers and transitions between minima on the protein’s free energy surface (FES). The complexity of the FES renders those large-scale fluctuations both anharmonic and anisotropic^2–5^. A common technique used to determine the slow, functional motions of proteins is the Principal Component Analysis (PCA)^3,5–8^. PCA is a dimensionality reduction procedure commonly used in signal analysis to highlight important persistent features underlying noisy data.^9^ When used to process atomistically-detailed molecular dynamics (MD) simulations, PCA reduces the dimensionality of the observed FES by identifying a few essential collective fluctuations ordered by decreasing eigenvalues.^10^ While PCA is computationally convenient, conceptually simple, and widely used, it lacks a physical basis beyond the empirical observations that it describes some slow, functional motions of a protein^7,11,12^.

In this study, we make a formal connection between the PCA expressed in the cartesian coordinates of a protein’s alpha-carbons and an approach we have developed to analyze the slow modes in protein dynamics, called the Langevin Equation for Protein Dynamics (LE4PD)^13–18^. The LE4PD theory, initially formalized as an isotropic equation of motion, is extended here to describe the anisotropic dynamics of proteins in the LE4PD-XYZ method. Like the PCA, the LE4PD-XYZ decomposes the protein’s motion into an orthogonal set of collective coordinates or modes, and, it captures the anisotropic, slow fluctuations of the protein, starting from the analysis of the atomistic MD trajectory. However, unlike the PCA, the LE4PD-XYZ is based on a Langevin equation of motion, which directly connects the large-scale fluctuations to the physical forces acting on the system. Because of its formulation, the LE4PD-XYZ allows for a detailed examination of the mode-dependent kinetics and fluctuation pathways.

Since the PCA is not directly related to an equation of motion, the amplitudes and timescales of fluctuations may be calculated by different procedures. For example, timescales have been calculated by integrating the autocorrelation function of the principal components.^4,19^ The direction and magnitude of the anisotropic fluctuations have been described using either the so-called ‘porcupine plots’^19–22^ or a simple linear interpolation between the extreme structures in the simulation trajectory^23^. In this manuscript, we compare the predictions of the PCA approach for the fluctuations with the largest amplitude to the results of the LE4PD theory applied to the same trajectory. By this analysis we quantify the importance of the different physical contributions to the PCA fluctuations, starting from the LE4PD-XYZ equation of motion.

The LE4PD-XYZ projects the simulation trajectory onto a mode-dependent free-energy surface. For each Langevin diffusive mode, our theory determines the three-dimensional energy landscape and the kinetic pathways of barrier crossing using a variant of the string method.^24,25^ Originally, the LE4PD measured the related kinetic times of barrier crossing by a simple Kramer’s rescaling of the friction coefficient, where we calculated the correction term from the height of the barrier, defined by the median absolute deviation (MAD)^16^. More recently, we paired the LE4PD with a Markov state model (MSM) analysis of the mode-dependent barrier crossing events. We then evaluated the mode-dependent kinetic times using the eigenspectrum of the slowest process predicted by the MSM analysis, and interpreted the results using the associated committor function.^18,26–28^ A similar analysis can be performed for the LE4PD-XYZ modes, and its comparison with the PCA modes is one of this study’s primary goals.

This study illustrates the advantages and limitations of the PCA normal mode decomposition compared with the Langevin formalism of the anisotropic LE4PD. PCA does not provide information on the timescales of the fluctuations. However, the LE4PD-XYZ equation, when hydrodynamic interactions are neglected, has forces acting between the amino acids that are directly related to the covariance matrix, and thus to PCA. The test system is a 1-*μs* MD trajectory of atomistic simulation of ubiquitin in an aqueous solution and physiological salt concentration. From its analysis we calculate the mode-dependent FES, its distinct pathways for protein fluctuations, and the related timescales using both the LE4PD-XYZ with and without hydrodynamics interactions. Then, we directly compare PCA’s linear fluctuations to the non-linear fluctuation pathway of LE4PD-XYZ. This analysis identifies the implications of neglecting hydrodynamic interactions and free energy barriers when PCA is extended to treat protein dynamics, by mapping the covariance matrix into the intramolecular matrix of the forces leading to the Langevin equation of motion.

## II. THEORY: THE LE4PD-XYZ EQUATION OF MOTION

In this section, we introduce the anisotropic Langevin equation for protein dynamics, or LE4PD-XYZ. The LE4PD-XYZ is a coarse-grained Langevin equation describing the fluctuations of the *i*^*th*^ alpha-carbon in a protein composed of *N* residues, and hence *N* alpha-carbons, from its equilibrium position,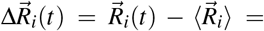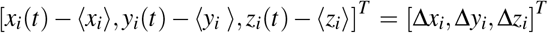. The equilibrium positions are defined as the time average over an MD trajectory consisting of *M* configuration points,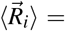 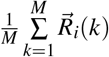,with *x_i_*(*t*), *y_i_*(*t*), *z_i_*(*t*) the distance of the *i*^*th*^ alpha-carbon from the origin of the simulation box at time *t* in the x−, y−, or z-direction, respectively. For a protein with *N* alpha-carbons there are a total of 3*N* degrees of freedom in the analysis, which is represented in the LE4PD-XYZ by the 3*N*-dimensional vector 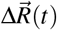:

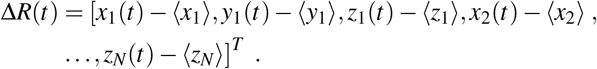

In the LE4PD-XYZ model, the time-evolution of a 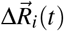 along the *α* direction,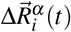, is given by the Langevin equation of motion

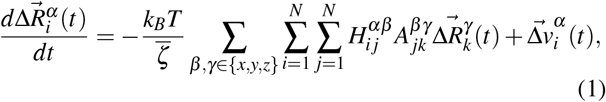

with *α, β, γ* ∈ *{x, y, z}* the coordinates in the three spatial dimensions. The equation is solved by applying the fluctuation-dissipation condition, as described in the Supplementary Material.

Here *k_B_* is the Boltzmann constant, T is the temperature in Kelvin, 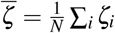 denotes the average residue friction coefficient, 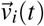 is a stochastic velocity, 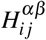 denotes the hydrodynamic interaction (HI) between the *α* component on bead *i* and the *β* component on bead *j* and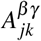 denotes the connectivity between the *β* component on residue *j* and the *γ* component on residue *k*. In Eq. 1, the hydrodynamic interaction matrix 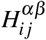 is given by

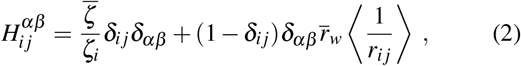

where *ζ_i_* the friction coefficient of residue *i*, 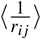 is the average inverse distance between residues *i* and *j*, and 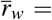 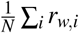 is the average residue radius exposed to the solvent.

The structural matrix, related to the mean-force potential, 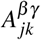 is defined as

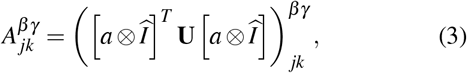

where 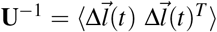 is the matrix of bond-bond correlations in Cartesian coordinates with 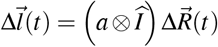, 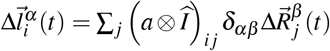, and *a* the *N* − 1 × *N* the incidence matrix that defines the connectivity between residues in the protein,

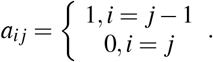

Here, *δ_αβ_* is the Kronecker delta, and the ‘⊗’ symbol denotes the Kronecker product.^29^ A detailed derivation of **H** as well as a formal connection of the **U** matrix defined here to the **U** in the previously developed, isotropic version of the LE4PD^13–16^ are given in the Supplementary Material.

It should be noted that the Langevin equation given in Eq.1 is identical in form to the optimized Rouse-Zimm equation for describing polymer dynamics derived by Zwanzig,^30^ excepting the detailed form of the **H** and **A** matrices, which here account for the chemical details of each residue and the semi-flexibility of the peptide bonds connecting the alpha-carbons.^13–16,18,31^ The Rouse-Zimm equation, without hydrodynamic interaction, can be derived from the Liouville equation, i.e. from the Hamiltonian of the system, by projecting the dynamics of the whole system onto the slow coordinates of the alpha carbons.^32,33^ The Rouse-Zimm equation is equivalent to a Fokker-Planck-Smoluchowski for polymer dynamics.^34,35^ In this respect, the LE4PD-XYZ equation presented here is founded on well-established first-principles approaches.

### A. Connecting the LE4PD-XYZ to PCA

The link between a PCA of the alpha-carbons and the LE4PD-XYZ method outlined above is as follows. For a given protein, an element of the covariance matrix in the Cartesian coordinates of the alpha-carbons is given by

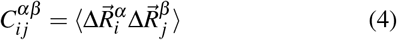

where 〈…〉 denotes, as above, an average over frames in the simulation trajectory. Using the definition of **A** given in Eq. 3 and that of **C** given in Eq. 4 it follows that

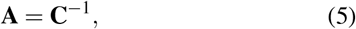

Thus, **A** and **C** possess the same set of eigenvectors and their eigenvalues are inverses of each other (provided **C** has full rank, which is always the case for a sufficiently long MD simulation).

The set of coupled Langevin equations, in Eq. 1, is diagonalized by the eigenvector **Q**, as **Q**^−1^**HAQ**= Λ, with Λ the diagonal matrix of eigenvalues of **HA**. By applying the eigenvector transformation, Eq.1 can be written in terms of its normal modes, 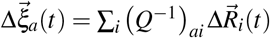 as

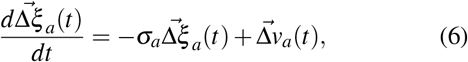

where 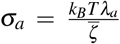 is the characteristic diffusive rate of mode *a*,^36^ with *λ_a_* = (Λ)*aa* the eigenvalue of mode *a*, and 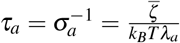 is the corresponding diffusive, barrier-free timescale of mode *a*. Finally, 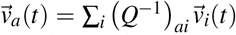 is the random velocity projected into mode coordinates. Note that 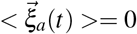, so 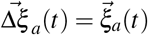.

Because of Eq. 5, it is straightforward to see that, when the hydrodynamic interaction matrix is approximated as an identity matrix, the PCA modes directly map onto the LE4PD-XYZ modes. Approximating the hydrodynamic interaction matrix in this way corresponds to assuming that i) the friction coefficient of each residue is set equal to the average friction coefficient, i.e.,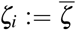, and that ii) the dynamical correlation due to hydrodynamics is neglected, i.e., **H**:= **I**, with **I** a 3N×3N identity matrix. Under those approximations, the eigenvalues (**Q**^−1^**AQ**)_*aa*_ = *μ_a_* where 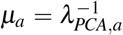 and *λ_PCA,a_* is the *a*^*th*^ eigenvalue of the covariance matrix. In this manuscript, we analyze the effect of these two approximations in the predicted timescale and amplitude of PCA fluctuations.

Furthermore, the LE4PD-XYZ method, even when neglecting hydrodynamic effects and simplifying the treatment of the residue-specific friction coefficients, accounts for the mode-dependent free energy barriers to transport in the mode space. This step is essential for an accurate description of the kinetics and transition mechanisms of the fluctuation dynamics in the mode coordinates,^14–16,18^ and is not part of the conventional PCA.

## III. METHODS

### A. Molecular Dynamics Simulations

We generated the MD simulation of ubiquitin using GRO-MACS version 5.0.4^37^ with the AMBER99SB-ILDN atomistic force field^38^ on the Comet supercomputer at the San Diego Supercomputing Center. The starting structure was selected from the Protein Databank, PDB ID: 1UBQ.^39^ We solvated the protein with spc/e water and minimized the energy using the steepest descent algorithm. We added Na^+^ and Cl^−^ ions until the ion concentration was 45 mM, with the concentration of ions selected to match the one used in nuclear magnetic resonance experiments of ubiquitin.^40^ Previously, we used those experimental data to test the accuracy of the LE4PD model against NMR data of T_1_, T_2_, and NOE relaxation experiments, with which the LE4PD approach show quantitative agreement.^14–16^ We subjected the protein-solvent system to two rounds of equilibration: first, a 50-ps equilibration in the NVT ensemble at 300 K, with a Nosé-Hoover thermostat controlling the temperature; then, a 450-ps NPT equilibration at 300 K, with the same thermostat and a Berendsen barostat set to 1 bar.

Following the NPT equilibration, we performed a 10-ns ‘burnout’ simulation at 300 K with the Nosé-Hoover thermostat to maintain the temperature constant. The last frame obtained in this procedure is adopted as initial configuration for the 1-*μ*s production run, which is performed using the same simulation parameters as the burnout simulation. Based on a manual inspection of the root-mean-squared deviation (RMSD) of the alpha-carbons from this first frame, we saw that the entire trajectory was fluctuating around an equilibrium value, and we used the entire 1-*μ*s of trajectory for the simulation analysis. We used the LINCS algorithm^41^ to constrain all hydrogen-to-heavy-atom bonds in the system and adopted an integration timestep of 2 fs during both the equilibration and the simulation run. The trajectory was recorded to file every 100 integration steps (every 0.2 ps), yielding a total of (10^6^ ps)/(0.2 ps/frame) = 5 × 10^6^ frames for analysis.

### B. Building the coarse-grained dynamical model of the anisoptropic Langevin equation, the LE4PD-XYZ

The LE4PD equation is a coarse-grained (CG) model for the dynamics of proteins. Each CG unit represents an entire amino acid in the protein’s primary sequence, with the center located at the position of the residue’s center-of-mass. The equilibrium configuration of the protein gives the equilibrium length of the connecting bonds between CG sites, while the site-specific friction coefficient of each amino acid, *ζ_i_* in Eq.2, is calculated from an extended Stoke’s law as^13^

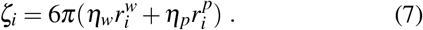

Here *η_w_* is the solvent’s viscosity, and *η_p_* is the viscosity in the hydrophobic core of the protein; 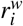 is the hydrodynamic radius of the amino acid for the solvent-exposed surface area, and 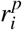 is the hydrodynamic radius calculated from the area exposed to the hydrophobic core. The internal viscosity *η_p_* is approximated as related to the water viscosity rescaled by the local energy barrier,^42^ *η_p_* = *η_w_* exp[<*E_int_* > /*k_B_T*] with <*E_int_* > ≈ *k_B_T* the minimal free energy barrier to the local internal motion of the protein.

Before performing the LE4PD-XYZ analysis, we processed the ‘raw’ MD trajectory to remove the rigid-body rotational and translational motions. This step is performed by first selecting the first frame of the simulation as the reference frame and then centering it at the simulation box’s origin. Concurrently, all the frames where the protein is broken across the periodic boundaries are made whole. Finally, all subsequent simulation frames are centered and superimposed to the reference structure by minimizing the mean-square difference between atomic positions. This procedure guarantees that six eigenvalues of **C** and **A**, which correspond to the rigid-body translational and rotational dynamics, are numerically indistinguishable from zero.

Because the spatial coordinates, which describe fluctuations around the mean value, have zero-mean and because the rigid-body rotational and translational motions of the protein are removed from the MD trajectory prior to analysis, the dynamics in the mode space are decomposed into a set of 3*N* −6 internal modes, plus 6 rigid-body modes corresponding to the rigid-body rotational and translational motions, which are associated with eigenvalues equal to zero.^43^ Because the protein conserves its globular shape during fluctuations, the removal of translation and rotation is a reasonable approximation, given that the coupling between rotation and fluctuations is minimized.^44^ The eigenvalues of the 3*N* −6 modes are ordered by ascending magnitude, *λ*_1_ < *λ*_2_ < *λ*_3_ <…< *λ*_3*N*_−_6_, with the smallest eigenvalues different from zero, corresponding to the largest amino acid fluctuation. Finally, since **C** is at least positive semi-definite, and **H** is also a positive definite matrix, both **A** and **HA** are at least positive semi-definite,^45^ which implies that *μ_a_*, *λ_a_* ≥ 0, ∀*a*.

### C. Validation of the LE4PD-XYZ theory

In Section II A we show how the eigenvalues of the PCA model relate to the eigenvalues and eigenvectors of the LE4PD-XYZ equation when hydrodynamic interactions and residue-specific friction coefficients are neglected and energy barriers are not included. Under those approximations, Eq. 2 is formally consistent with a Fokker-Planck-Smoluchowski equation, where the dynamics are expressed as a function of the probability density function.^34,46^ In reference^35^, Hinsen et al. analyze the anharmonicity of protein fluctuations starting from the Fokker-Planck-Smoluchowski equation, and by modeling with this equation the time decay of the density fluctuations as measured in neutron scattering experiments. The theoretical predictions in that study are consistent with the time decay observed in their simulations. The parameters entering the equation of motion were obtained by direct comparison with the simulation trajectories. The length of the simulation is limited to 1.5 ns, during which crossing of large energy barriers is unlikely to occur.

Figure 1 shows a direct comparison between the predictions of the LE4PD-XYZ for the time decay of the residue fluctuations and the same properties directly calculated from the trajectory. The parameters entering the LE4PD-XYZ equation are the amino acid friction coefficients and the energy barriers in the mode representation, which are calculated as described in Sections III B and IV. The mode-dependent timescale, which defines the decay of the local fluctuations, is simply rescaled using Kramers’ theory of reaction kinetics and the height of the energy barrier (see Section IV for details).^47^ The agreement is close to quantitative for the time correlation functions (tcfs) shown in Figure 1. The observed good agreement between theory and simulations depends on including the mode-dependent energy barriers and the hydrodynamic interactions. This observation may seem to be at odds with the good agreement observed in reference^35^ between simulations and the Fokker-Planck-Smoluchowski equation, where hydrodynamics and energy barriers are not included. However, in reference^35^ the simulation is limited in length to 1.5 ns, while our study simulations are 1 *μs* long, and during that time several crossing of high energy barriers may occur.

**FIG. 1.**
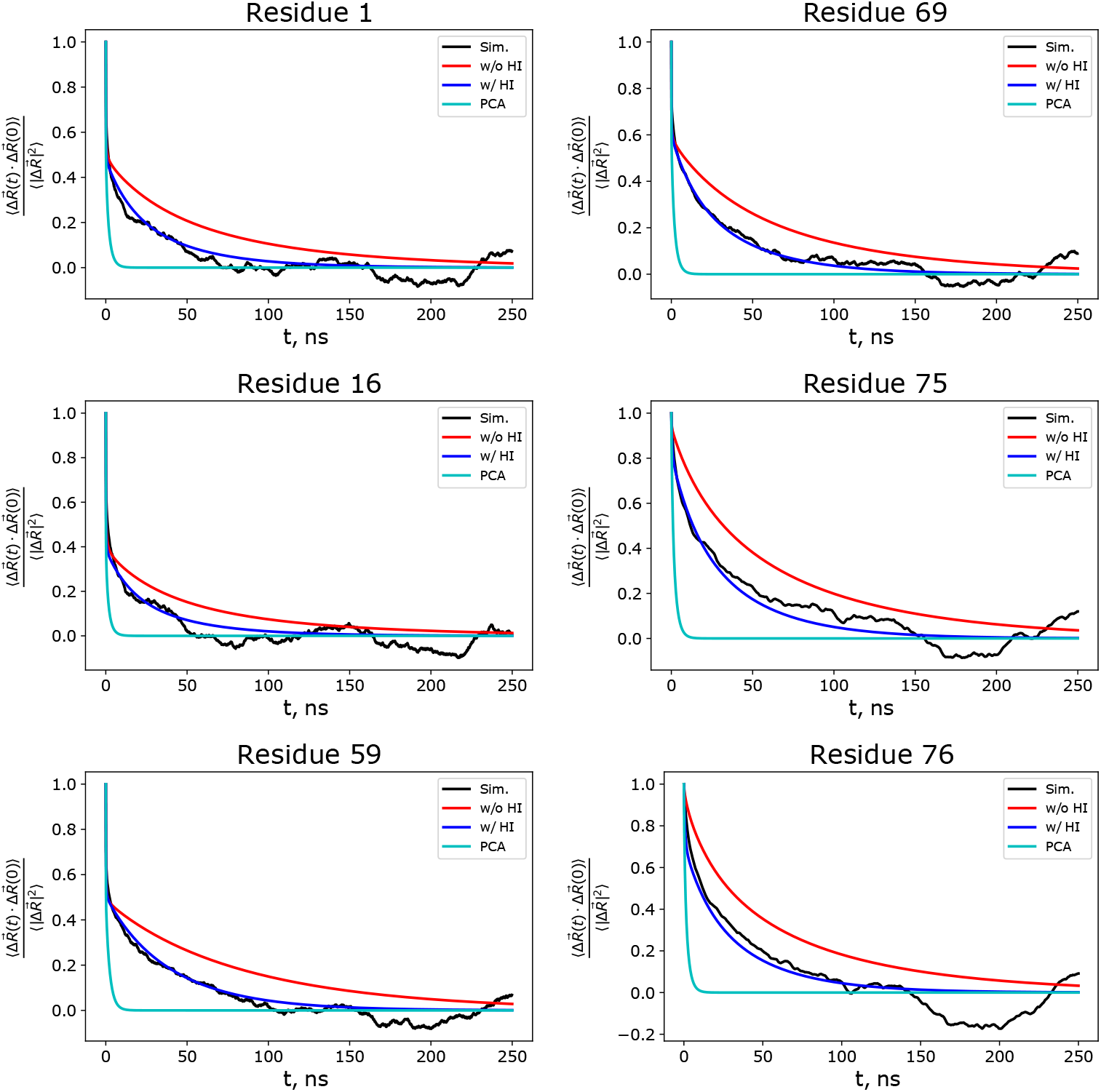
Time decay of the position fluctuation for different residues in the ubiquitin. The predictions of the LE4PD-XYZ theory with hydrodynamic interactions included (blue) are compared with the theoretical predictions without hydrodynamics (red) and with simulations (black). The predictions of the LE4PD-XYZ theory when hydrodynamic interactions and the mode-dependent internal energy barriers are both excluded (cyan) correspond to the decay of the fluctuations given by the PCA eigenvalues. The agreement between the LE4PD-XYZ data, when energy barriers and hydrodynamic interactions are included, is close to quantitative.

In Figure 1 we report as an example the decay of the fluctuations for six individual amino acids along the protein primary structure. The figure shows the LE4PD-XYZ predictions, with and without hydrodynamic interactions, while the correction due to the internal energy barriers is included. The agreement between theory and experiments is close to quantitative. It also shows the predictions of the LE4PD-XYZ theory without hydrodynamic interactions and without the correction due to the mode-dependent internal energy barriers; those are the predictions obtained if one calculates the timescale directly from the eigenvalues of the covariance matrix, i.e. from PCA. Including hydrodynamics interactions and energy barriers when modeling the dynamics of polymers is important for the good agreement with the data. The inclusion or exclusion of the HI contribution modifies the final decay of the tcfs. In fact, hydrodynamic ensures the correct scaling exponent with time of the polymer’s long-time dynamics as observed experimentally, for example, in neutron scattering.^34^

## IV. CONTRIBUTION OF THE HIGH ENERGY BARRIERS IN THE FLUCTUATION DYNAMICS OF PROTEINS DETECTED BY THE PCA AND LE4PD-XYZ METHODS

The Langevin equation expresses the time evolution of the protein’s motion by identifying the forces that act on each amino acid. Those forces define how the protein’s dynamics evolve in time and include forces between amino acids due to the intramolecular potential of mean force and long-range interactions mediated by the solvent. When the Langevin equation represents the dynamics of a protein in solution, the solvent’s effect enters through a residue-dependent friction coefficient and the hydrodynamic interaction matrix (see Eq. 2).

When the hydrodynamic interaction matrix reduces to an identity matrix because residue-specific friction coefficients and long-range forces mediated by the solvent are neglected, the time evolution of the protein’s motion follows the covariance matrix. The latter describes harmonic fluctuations of the amino acids away from their equilibrium position. Under this approximation, the covariance matrix’s eigenvalues are the inverse of the ones from the LE4PD-XYZ approach, and define identical timescales of the dynamics. Thus one may conclude that the dynamics described by the PCA’s eigenvalues follows a simplified protein’s equation of motion, which neglects the specificity of the amino acid friction coefficient and the solvent-mediated amino acid interactions.

The first approximation that one needs to enforce to recover the PCA dynamics from the Langevin equation is the assumption that all the amino acids in the protein have identical friction coefficients. The friction coefficient is proportional to the hydrodynamic radius of the residue, as discussed in Section III B. This radius may vary with the residue’s chemical nature and its location inside the protein’s three-dimensional structure. It depends on the extent of the surface exposed to the solvent, which can change dramatically for different amino acids along the protein’s primary sequence.^48,49^ Thus, there is no physical motivation to support the adoption of an identical friction coefficient for all the amino acids in a protein. We show how this approximation affects the dynamics in Section V.

The second approximation assumes neglecting the hydrodynamic interaction. The hydrodynamic interaction matrix represents the long-ranged interactions between amino acids, mediated by the solvent described as a continuum medium.50 When describing the rotational and translational dynamics of proteins, it is common practice to include hydrodynamics effects. However, hydrodynamic contributions to proteins’ internal motion are, sometimes, neglected. While this may be a reasonable approximation for very localized motion, such as local vibrations, in general, hydrodynamic effects are not negligible. The non-local hydrodynamic coupling of amino acids’ dynamics can alter the time scale of the large-amplitude fluctuations. We will study in more detail the effect of hydrodynamic interactions on ubiquitin’s mode fluctuations in Section V.

### A. Building Free Energy Surfaces

An important contribution to the timescale of protein’s fluctuations is the crossing of high energy barriers in the FES. Note that this contribution is present even when the hydrodynamic interaction is neglected. The Langevin equation, given in Eq.1, is a diffusive approach that does not explicitly account for energy barriers along the mode coordinates 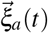 and, like PCA, describes harmonic fluctuations away from the equilibrium structure. In the absence of hydrodynamics, where the fluctuations in PCA and LE4PD-XYZ are driven by the same covariance matrix, our approach allows one to calculate for each mode, *a*, the associated free-energy map. From the free energy surface it is possible to quantify the barriers to transport, thus obtaining an accurate determination of the kinetics of the conformational fluctuations in the mode coordinates^16,18^ (see Section IV A).

We calculate the mode-dependent FES from the MD trajectory by projecting the position vectors into the modes, using the LE4PD-XYZ eigenvectors. In the absence of hydrodynamic interactions, these are also the PCA eigenvectors. In the three-dimensional description, it is convenient to write each eigenvector as the sum of its components along the x-, y-, and z-directions

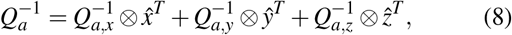

with 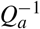 the *a^th^* row of the **Q**^−1^, which is the matrix of the left eigenvectors of the product **HA**. Here 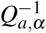, with *α* ∈ {*x, y, z*} is an element of the 3*N* × *N* matrix describing the projection of energy barrier calculated for LE4PD-XYZ mode *a* using the median absolute deviation (MAD)52,53 from the minimum of the x-, y-, and z-coordinates of each alpha-carbon onto mode *a*, while 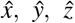 are the basis vectors in the x-, y-, and z-directions, respectively, e.g., 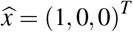. The projection of the simulation trajectory using the eigenvector matrix defined above leads to the mode coordinates along the three spatial directions

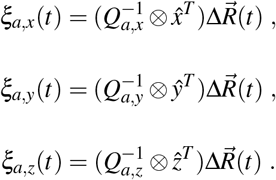

From these mode vectors, it is possible to build an FES and more easily visualize the FES by calculating the probability in spherical coordinates. With the mode definition in hand, the polar and azimuthal angles of 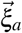 are

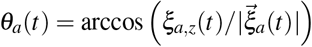

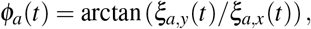

with 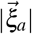 the magnitude of 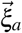. For each mode, we derive the FES by binning into a 2D-histogram the probability of occupying a given value of *θ_a_* and *ϕ_a_*, 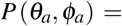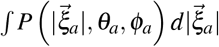 and then performing a Boltzmann inversion of the probability. The probability distribution in all theoretical calculations is a function of the three spherical coordinates. However, the graphical representation of the free energy surface is simplified when omitting the probability as a function of 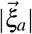, as the energy plot reduces to three dimensions. This step is possible because the free energy as a function of 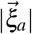 does not present any remarkable feature. Thus, the total probability can be averaged over the values of 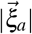 without losing important information. Instead, distinct dynamical pathways are visible in the FES as a function of the polar and azimuthal angles. Thus, the mode-dependent free energy surface is conveniently described by the free energy as a function of the *θ_a_* and *ϕ_a_* angles, while we average over the surface’s fourth dimension, 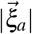. The free-energy for a given (*θ_a_, ϕ_a_*) pair, as sampled by mode *a*, reduces to

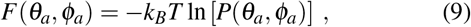

averaged over all the values of 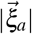. A further discussion of this step is presented in the Supplementary Material.

To account for the effects of energy barriers in the decay of the correlations of the residue fluctuations, such as those shown in Figure 1, the friction coefficient in Eq. 6 is re-normalized using a Kramers-type approach^17,51^: 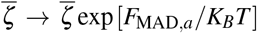, where *F*_*MAD,a*_ = median(|*F* (*θ_a_, ϕ_a_*) − min(*F* (*θ_a_, ϕ_a_*)) |) is an average free-energy barrier calculated for LE4PD-XYZ mode *a* using the median absolute deviation (MAD)^52,53^ from the minimum of energy on the surface. This approach rescales the diffusive mode timescales as 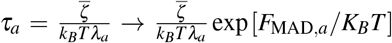 Using the MAD statistic removes any poorly sampled regions of the energy surface when the calculating the barrier, and using the MAD to rescale the friction coefficients has previously been shown to be effective in describing the slow-down in the decay of the M_1_(t) time correlation function calculated from the LE4PD theory at the 2.5 ns timescale for ubiquitin,^14^ the same protein under study here.

We obtain a free energy map for each mode, which presents a complex landscape, with minima, maxima, and complex dynamical pathways (see for example Figure 2). In the following, we will study the energetic pathways that emerge from the LE4PD-XYZ when hydrodynamic interactions are neglected. As mentioned earlier, the LE4PD-XYZ directly connects with PCA in the ‘free-draining’ limit, where Eq.1 is solved while assuming **H** ≔ **I**. In that case, the mode solutions and the free energy maps from both approaches are identical.

**FIG. 2.**
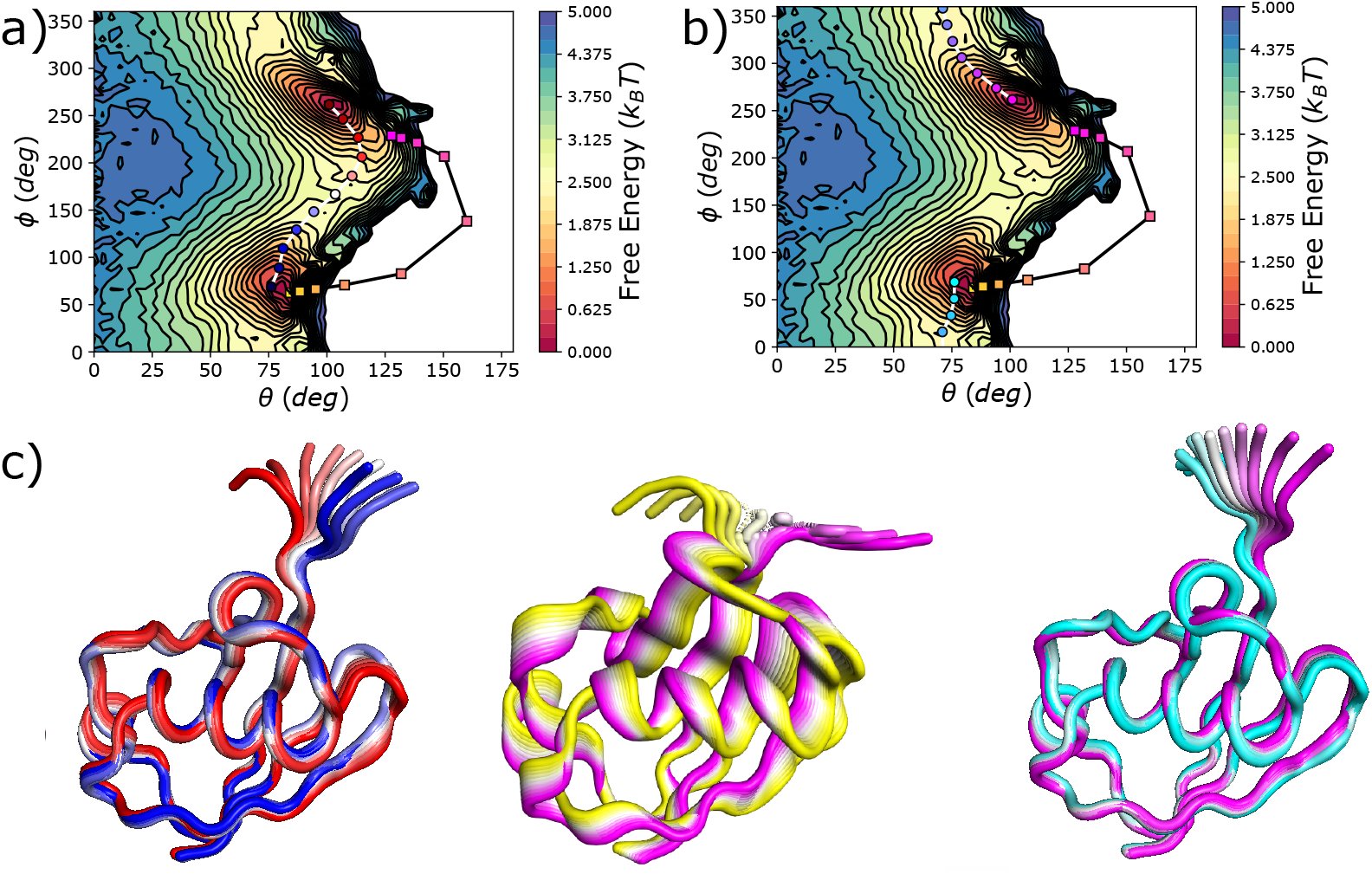
Free-energy surface for the first LE4PD-XYZ mode, solved for the case where **H**:= **I**, so that the LE4PD-XYZ mode solutions are identical to the calculated PCA modes. The FES in the a) and b) panes are identical. a) The blue-white-red path is a minimum energy pathway between the two minima, found using the string method. The yellow-white-magenta path shows the PCA trajectory of the linear interpolation between the two extreme structures. b) The cyan-white-magenta path follows a second minimum energy pathway that crosses the periodic boundary at *φ* = 0 (*deg*) = 360 (*deg*). The PCA linear-interpolation trajectory is identical to the one in the a) panel. c) The real-space, 3D fluctuations corresponding to each of these pathways are depicted by the superimposed ubiquitin structures. Each structure is colored corresponding to the analogously colored image along the pathway.

### B. Comparing Pathways through the Free-Energy Surface

Figure 2 illustrates the free energy surface (FES) and the kinetic pathways of the first LE4PD-XYZ/PCA mode, which corresponds to the first non-zero eigenvalue. In the top two panels (a) and (b) the FES is identical, while the pathways are different. The FES presents two well-defined minima in energy, one at *θ_a_* ≈ 75 (*deg*) and *ϕ_a_* ≈ 60 (*deg*) and the second at *θ_a_* ≈ 100 (*deg*) and *ϕ_a_* ≈ 270 (*deg*). Superimposed to the FES are the kinetic pathways of crossing the free energy barriers. Because the angular coordinate *ϕ_a_*, is periodic, there are two possible pathways to connect the two minima in the free energy surface. Those are calculated using a variant of the string method,^18,24^ and are reported in the Figure 2a and Figure 2b, top panels. For the PCA, instead, the path is defined as the linear interpolation between the most extreme configurations sampled by the simulation^23^ and is identical in the two panels. Interestingly, these paths crossing the energy barriers and the PCA linear interpolation do not coincide.

The PCA interpolation method gives a pathway that does not quite begin or end in the minima of the free-energy surface. The path’s extrema capture some less-likely configurations populated by the fluctuations of the mode around the most probable configurations. We also observe that the PCA linear interpolation does not follow the pathway of the energetically-favored barrier crossing. The intermediate states cross through a low-probability region of the surface and do not travel through the ‘valley’ between the two minima.

Figure 2c presents the superimposed configurations that populate the three kinetic paths just mentioned. The real-space structures that the molecule experiences while moving along the two most probable paths of barrier crossing in the FES are depicted in the left and right panels. The central panel, instead, shows the set of conformations populating the interpolation path of PCA. The PCA structures, in the central panel of Figure 2c, show large deformations along the entire alpha-carbon sequence of ubiquitin. Instead, the two pathways predicted by the LE4PD-XYZ show that the motion is concentrated in the tail region of ubiquitin.

A reason for the inconsistency with LE4PD of the interpolation method is its reliance on the extreme values of *ξ_a_*(*t*) = (**Q**^−1^Δ*R*)_*a*_(*t*). Thus, it utilizes only two samples of *ξ_a_*(*t*) to generate a fictitious trajectory for visualization, while the pathway method used in the LE4PD-XYZ utilizes the entire *ξ_a_*(*t*) trajectory to create the FES and hence the pathways between minima on the FES. And while minima on the FES and extreme values of *ξ_a_*(*t*) tend to be correlated, the extrema of *ξ_a_*(*t*) are by no means guaranteed to represent the configurations of the energetic minima. For example, Figure 3 shows the projection of *ξ_a_*(*t*) onto the FES’s for LE4PD-XYZ modes 1 and 7 without HI. While the lowest and highest values of *ξ_a_*(*t*) are situated in the FES’s deepest minima, the absolute minimum and maximum of *ξ_a_*(*t*) may not be of the lowest energy. If they were, the locations marked by the colored stars (maximum and minimum projections of *ξ_a_*(*t*)) and triangles (minima of energy) would superimpose in both cases. Figure 3 also shows that the most extreme projections of *ξ_a_*(*t*) tend to lie outside the deepest minima of the FES. The displacement can be small, as in the extreme positive projection of *ξ_a_*(*t*) for the first mode and in both the extreme projections for mode 7, or it can be different by a more substantial amount, as is the case for the extreme negative projection of mode 1. The minima in the FES and the extrema of the PCA fluctuations may coincide. However, even in this case, there is no guarantee that the intermediate states in PCA, found using the interpolation procedure, define an energetically favorable pathway.

**FIG. 3.**
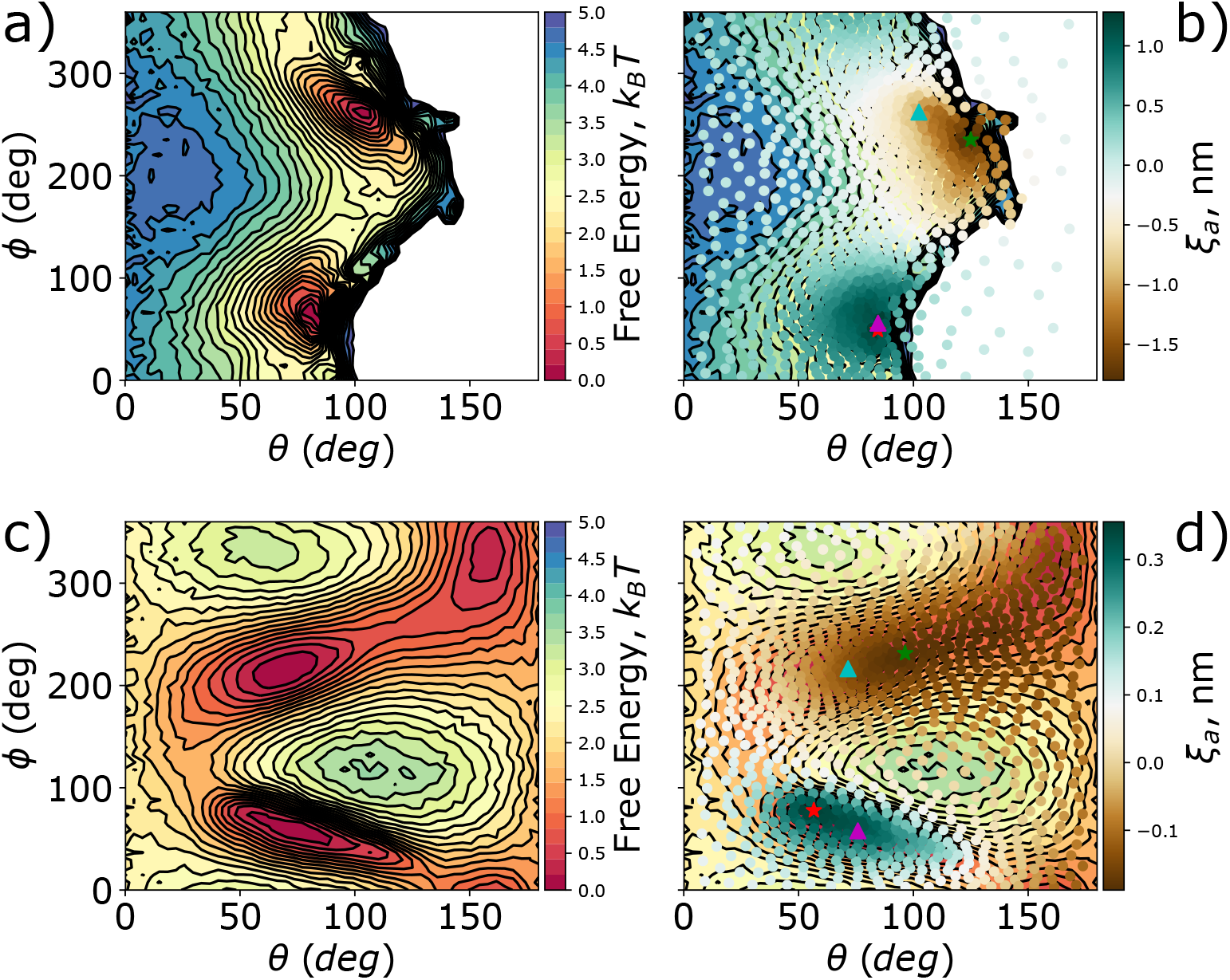
a) FES for the first internal LE4PD-XYZ mode. b) Projection of *ξ_a_* (green-white-brown markers) onto the two-dimensional FES for a=1. c) FES for the seventh internal LE4PD-XYZ mode. d) Projection of *ξ_a_* (green-white-brown markers) onto the two-dimensional FES for a=7. Green and red stars mark the locations with the lowest and highest projection of *ξ_a_*, respectively. Cyan and magenta triangles mark the locations with the lowest free-energy, subject to the constraint of having either a negative or positive projection along *ξ_a_*, respectively.

## V. HOW INCLUDING HYDRODYNAMICS MODIFIES EIGENVALUES, EIGENVECTORS AND RELATED QUANTITIES: A COMPARISON OF PCA AND THE DIFFUSIVE LANGEVIN APPROACH OF THE LE4PD-XYZ

The study reported in the previous section shows that the LE4PD-XYZ approach identically maps onto the PCA formalism when the hydrodynamic interaction is neglected because the LE4PD-XYZ eigenvalues and eigenvectors map directly onto the PCA eigenvalues and eigenvectors. It also shows that the simple interpolation procedure of PCA portrays an approximate representation of the slow motion, as the path of the fluctuation may not follow the kinetic pathway of minimum energy between two energetic minima. Thus, the PCA’s amplitude and pathway of slow fluctuations can be somewhat inaccurate in representing the most-probable and, likely, biologically-relevant fluctuations in the protein.

Modifying the forces acting on the protein by including hydrodynamics interactions modifies the eigenvalues and eigen-vectors of the **HA** matrix product. This may change the timescale and amplitudes of the slow fluctuations around the equilibrium configuration. As a first step, we focus on comparing eigenvalues and eigenvectors with and without hydrodynamic interactions. Note that when we include hydrodynamic interactions, we also assume residue-dependent friction coefficients, which are calculated following the procedure described in the Methods section.

Figure 4 shows how the eigenvalues are modified by the inclusion of residue-specific friction coefficients in the hydrodynamic interaction matrix, and by the inclusion of the full HI matrix with long-ranged cross interactions and residue-specific friction coefficients. Given that the mode-dependent timescales are defined as the inverse of the eigenvalues of the matrix product **HA**, 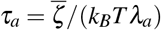, one can see that including the HI ‘flattens’ the eigenvalues, decreasing the timescale of the lowest-index modes, making them faster, and increasing the timescale of the highest-index modes, making them slower. The theory of polymer dynamics predicts a similar effect: the scaling exponent of the Rouse modes is modified by the inclusion of the hydrodynamic interaction leading to the Rouse-Zimm approach, from which the LE4PD is derived. Including hydrodynamic effects ‘softens’ the eigenvalue spectrum by lowering the magnitude of the dynamic scaling exponent^15,34^.

**FIG. 4.**
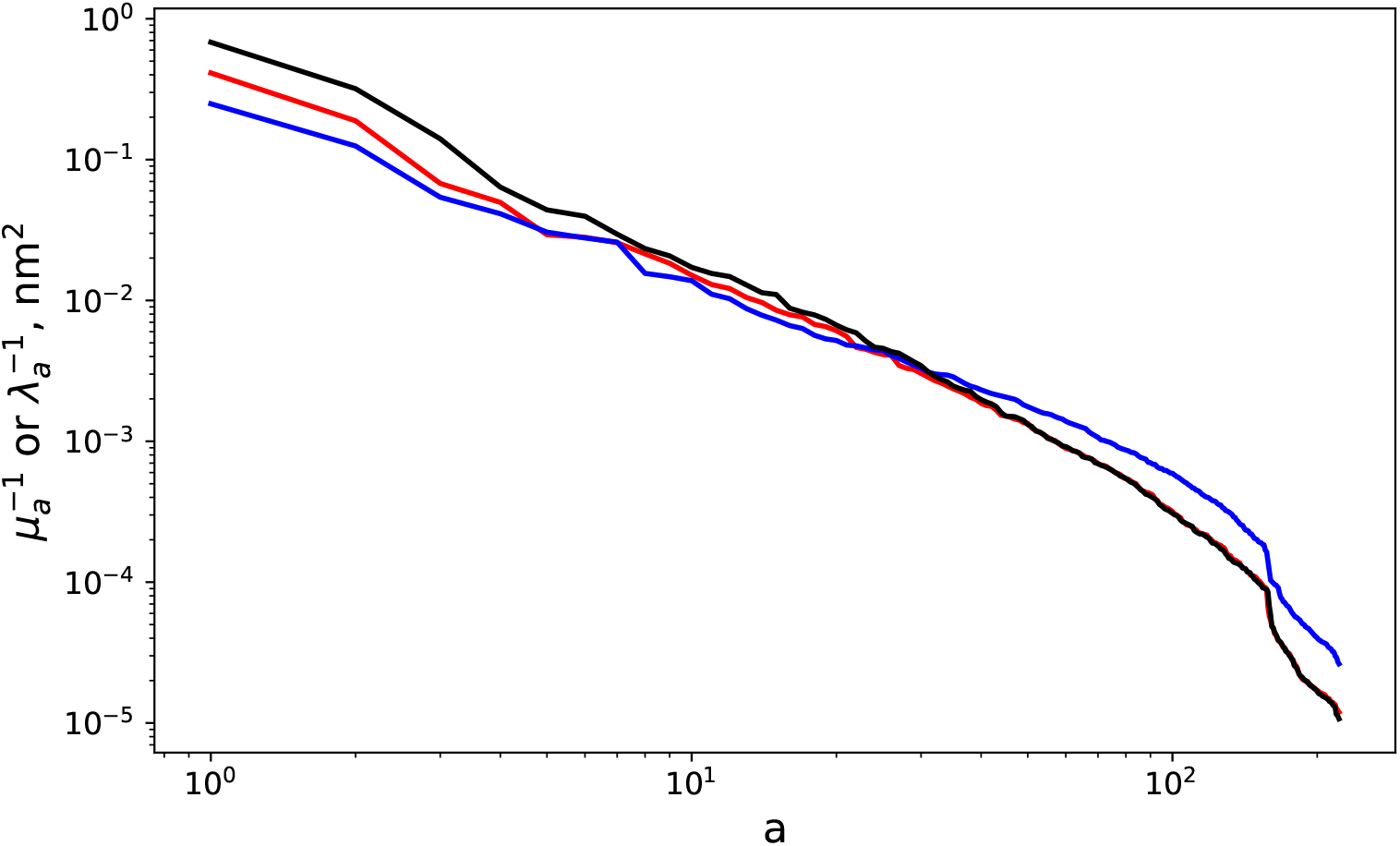
Comparison of the eigenvalue spectrum without hydrodynamic interaction (black curve), with the ones where the reside-specific friction coefficients are included, to account for the chemical specificity of each residue (red curve), and with the eigenvalues resulting from the diagonalization of the product of matrices containing the full hydrodynamic interaction matrix (blue curve).

Modifying the description of the hydrodynamic interaction is likely to affect the values of the eigenvectors of the matrix product **HA** as well. Figure 5 compares the eigenvector projections into the x-, y-, and z-coordinates, with and without HI, for the five slowest LE4PD-XYZ modes. Note that the timescales of these modes are given by the inverse of the LE4PD eigenvalues: including the energy barriers may modify the modes’ order.

**FIG 5.**
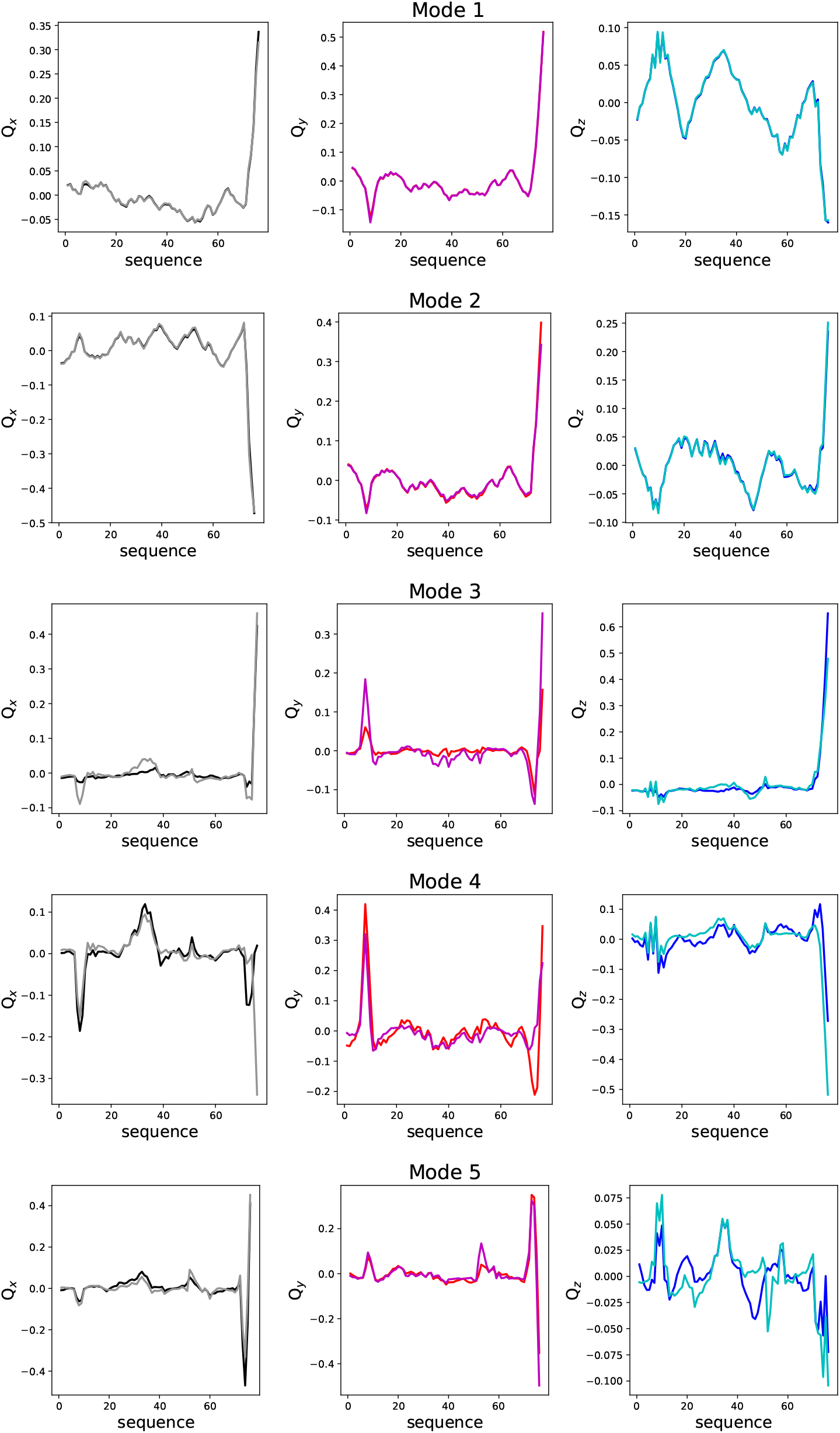
Comparison of the 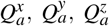, without (black, red, blue) and with (grey, magenta, cyan) hydrodynamic interactions for the first 5 LE4PD-XYZ modes.

For the three slowest modes, the eigenvectors are essentially indistinguishable whether hydrodynamic interactions are included or not; however, differences become more apparent for modes 4 and 5. Because for a given mode the eigen-vector determines the position and the amplitude of the fluctuations along the primary sequence of the protein, we expect that the inclusion of hydrodynamics will not affect the location of the slow fluctuations, but may modify their amplitude.

The direct comparison of the eigenvectors may be affected by the different ordering of the eigenvalues in the complete (with HI) and approximated (without HI) formalism, because the ordering of the eigenvectors depends on the ordering of the eigenvalues. Including the HI can modify the frequency of some mode fluctuations, thus changing the ordering of those modes. For example, the fluctuation of a loop could become slower due to the presence of long-ranged interactions once the hydrodynamics is included. To study possible cross-correlation between modes of different number, we calculated the overlap matrix, *O_ab_*, between the eigenvector of mode *a* calculated without hydrodynamics (i.e. in PCA, *Q_a_*), and the eigenvector of mode *b* calculated with HI included,(i.e. in LE4PD-XYZ, 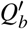), as

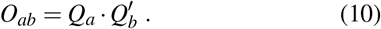

Since each eigenvector is normalized, the overlap matrix has the dimension of the number of internal modes, squared. For ubiquitin, a protein composed of 76 residues, 3N − 6 = 222 and *O* is a 222 × 222 matrix. A more in-depth explanation of the significance of *O* is given in the Supplementary Material.

Figure 6 shows *O*, which, overall, has a weakly diagonal structure, indicating that there is a similarity between the dynamical processes described by both types of LE4PD-XYZ treatments. However, the trace of *O* is only 34.8, while a perfect correspondence between processes would yield a trace of 3N − 6 = 222. For the ten slowest modes, which are represented by the insert in the up right corner of Figure 6, the overlap between eigenvectors from the two treatments is large, especially for the three slowest modes and the fifth slowest mode, each of which possesses an overlap greater than 0.91. Furthermore, from the trace of the overlap matrix we can calculate the average overlap of the first ten modes, which is 0.75. Subtracting this contribution from the total trace of 34.8, the remaining 212 internal modes have an average overlap of 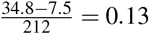, which is small, indicating little similarity between the higher-index, fast modes from the two approaches. Thus, including the hydrodynamic interactions in the equation of motion produces a significant alteration of the highest index modes, which are fast and have small amplitude. The changes of the eigenvectors corresponding to the slowest modes appear, instead, to be more contained, at least for the protein ubiquitin, which is the focus of this study. These results suggest that we should expect small changes in the FESs corresponding to the slowest dynamical modes when hydrodynamic interactions are included.

**FIG. 6.**
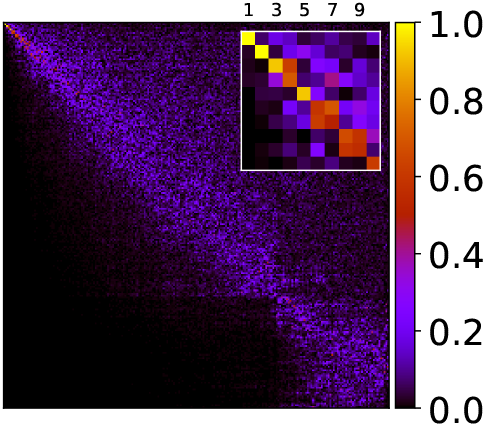
Overlap matrix *O*, defined in Eq. 10, between the right eigen-vectors of the LE4PD-XYZ without hydrodynamics and the right eigenvectors of the LE4PD-XYZ with hydrodynamic interactions included. Overall, there is weak diagonal trend in this matrix, indicating similarity between the analogous modes in both approaches. Inset: Sub-matrix of *O* corresponding to the overlap between the first ten modes from each LE4PD-XYZ treatment. Scale bar for the overlap between each mode is given to the right of the plot.

## VI. EFFECT OF INCLUDING HYDRODYNAMIC INTERACTIONS ON THE POSITION AND AMPLITUDE OF SLOW MODE FLUCTUATIONS: A COMPARISON OF PCA VERSUS THE DIFFUSIVE LANGEVIN APPROACH OF THE LE4PD-XYZ

The previous section of this paper has shown that there are small but significant differences emerging in the eigenvalues and eigenvectors when one includes hydrodynamic interactions in the equation of motion. Thus, the eigenvectors used to map the simulation trajectory onto the normal modes and build the FES in the PCA are different from the ones in the anisotropic LE4PD with hydrodynamic interaction. So, it is likely that including the hydrodynamic interaction will lead to timescales and location of fluctuations that are different from the ones measured in PCA. In general, one may expect the timescales of processes that include hydrodynamic interactions to be more realistic because their dynamics follow an equation of motion that properly accounts for the effects of the solvent.

However, in the case of ubiquitin, the eigenvectors of the first two modes are almost quantitatively identical (see Figure 5) while the corresponding eigenvalues are modified by the presence of hydrodynamic interactions (see Figure 4). Thus, we expect the free energy maps for those two modes to be very similar in PCA and LE4PD-XYZ, while the timescale of fluctuations may be different.

The FES is calculated by mapping the mode 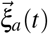 in polar coordinates, where 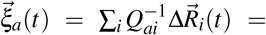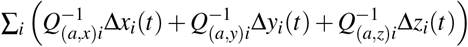. The resulting free energy surfaces, *F*(*θ_a_, ϕ_a_*), of the first five LE4PD-XYZ modes are displayed in Figures 7 and 8. The ordering of the modes in Figures 7 and 8 is based on the eigenvalues of the **HA** matrix, which do not include the slow-down in the mode-dependent dynamics due to the inclusion of free-energy surfaces.

**FIG. 7.**
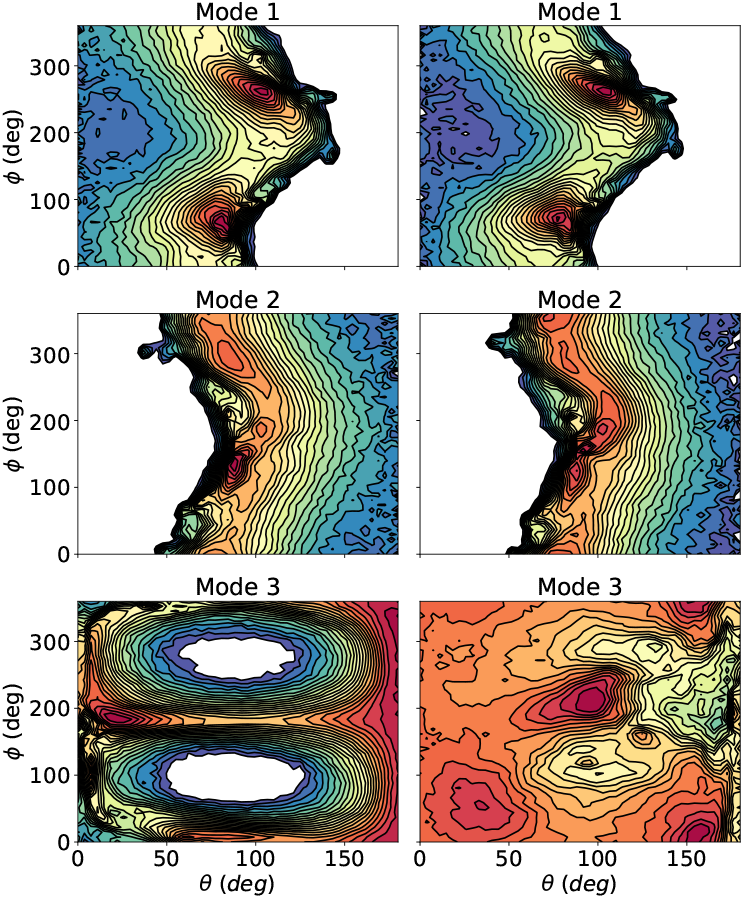
Comparing the three slowest modes without (left) and with (right) hydrodynamics in the LE4PD-XYZ analysis. Red corresponds to low energy and blue to high energy; all regions with a free-energy above 5 k_*B*_T are ‘masked’ as white. The scaling of the free energy is the same as that in Figures 2 and 3.

**FIG. 8.**
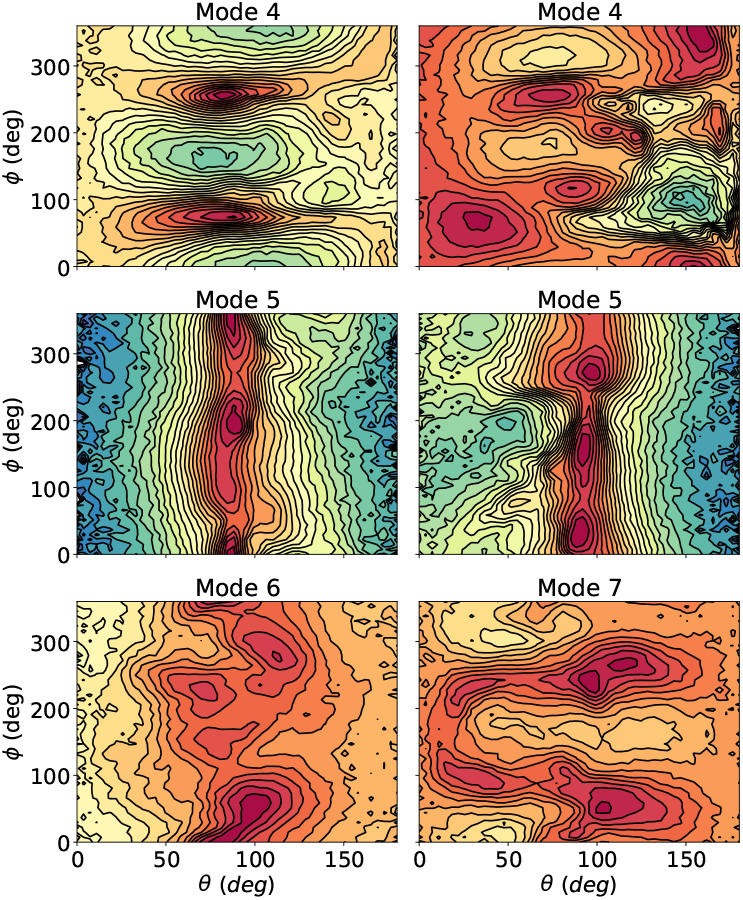
Comparing the three next slowest modes without (left) and with (right) hydrodynamics in the LE4PD-XYZ analysis. Red corresponds to low energy and blue to high energy; all regions with a free-energy above 5 k_*B*_T are ‘masked’ as white. Mode 7 is swapped with mode 6 in the hydrodynamics treatment because it overlaps more strongly with mode 6 without hydrodynamics.

Each mode with HI included is compared directly to the one without, considering modes that have the highest overlap according to Eq. 10. In agreement with Figure 5, the first two modes and the fifth mode in both approaches have striking similar free-energy surfaces. In contrast, modes 3, 4, and 6 or 7 are quite different, which is also in agreement with the mode-mode overlap matrix *O* shown in Figure 6. Note that mode 6 with HI (LE4PD-XYZ) has the maximum overlap with mode 7 without HI (PCA), and they will be directly compared.

To analyze the position and amplitude of the local fluctuations along the primary sequence of ubiquitin, we calculate the total mean-squared fluctuations of the alpha-carbons, as follows:

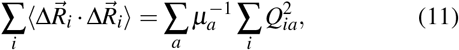

By isolating the element in the first sum corresponding to mode *a* and the element *i* in the second sum corresponding to residue *i* in the protein on the right-hand side of Eq. 11 we obtain the definition of the mean-square fluctuations at residue *i* due to the process described by mode *a*, which we will call the mean-squared local mode lengthscale, 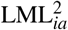:

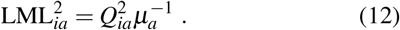

In the anisotropic formalism of LE4PD-XYZ, 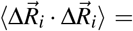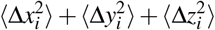, and, by partitioning the *Q_a_* into its *x*−, *y*−, and *z*−components, the 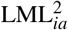 can be decomposed into *x*−, *y*−, and *z*−projections:

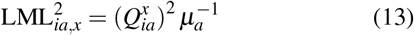

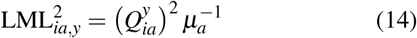

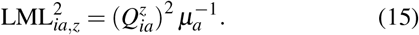

The use of the anisotropic 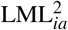 provides information on location, amplitude, and directionality of the localized fluctuations. The LML for the six slowest LE4PD-XYZ modes are shown in Figures 9 and 10; as previously, the modes with the highest overlap in LE4PD and PCA are compared directly in each of the subplots of the figures. Qualitatively, in most of the slowest modes, more specifically in the first three modes, there is little difference between the mode-dependent *location* of the fluctuations predicted whether hydrodynamic effects are included or neglected, in agreement with what we observed in the eigenvectors (see Figure 5). For these slow modes, the anisotropy of the fluctuations is not changed; the fluctuations in the *x−, y−,* and *z*–coordinates are the same regardless of the level of theory chosen, PCA or LE4PD-XYZ.

**FIG. 9.**
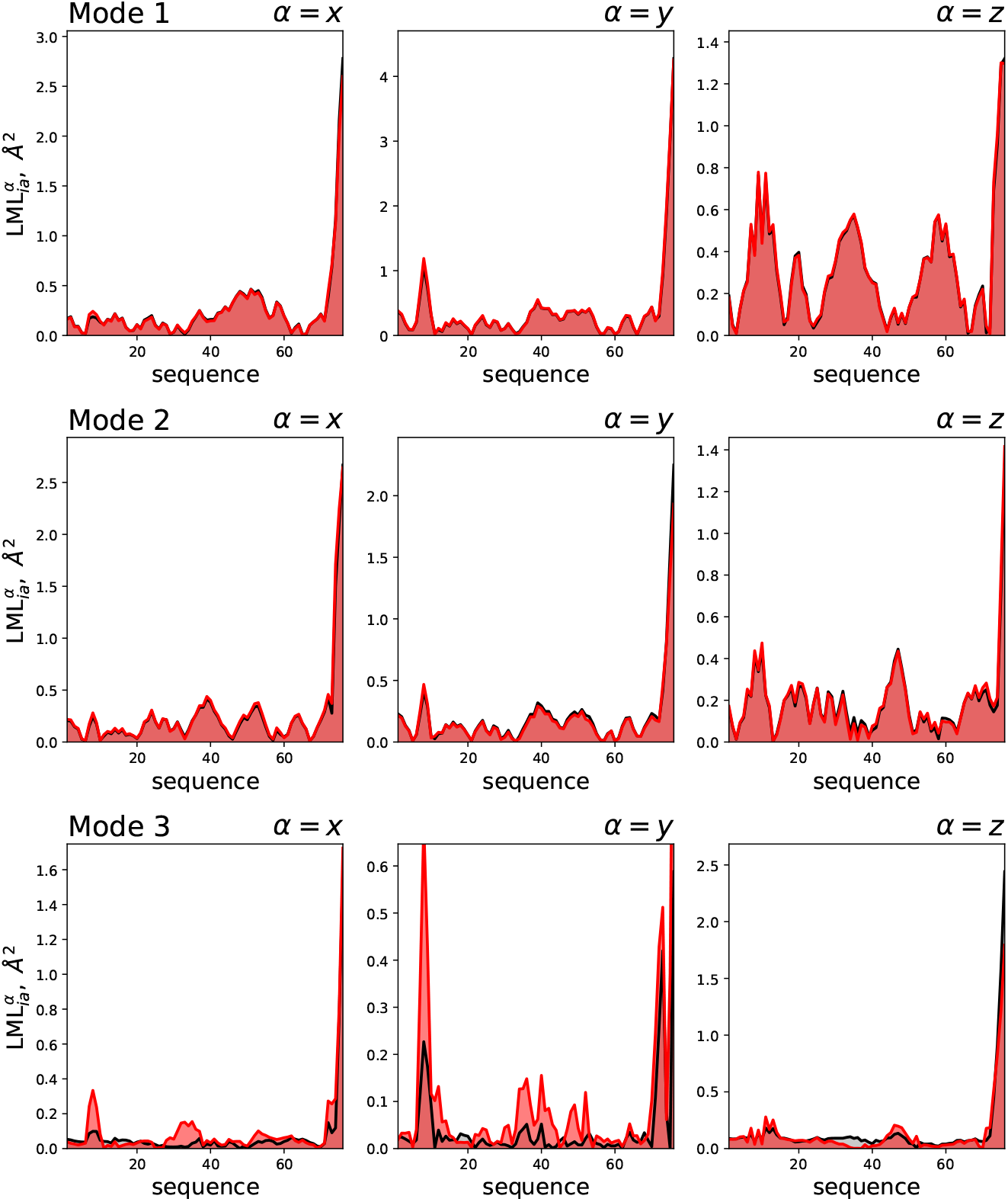
Anisotropic LML, 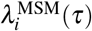 for the first three LE4PD-XYZ modes, as ordered by the *λ_a_* eigenvalues, for the case where hydrodynamic effects are neglected (black) and with hydrodynamic effects included (red).

**FIG. 10.**
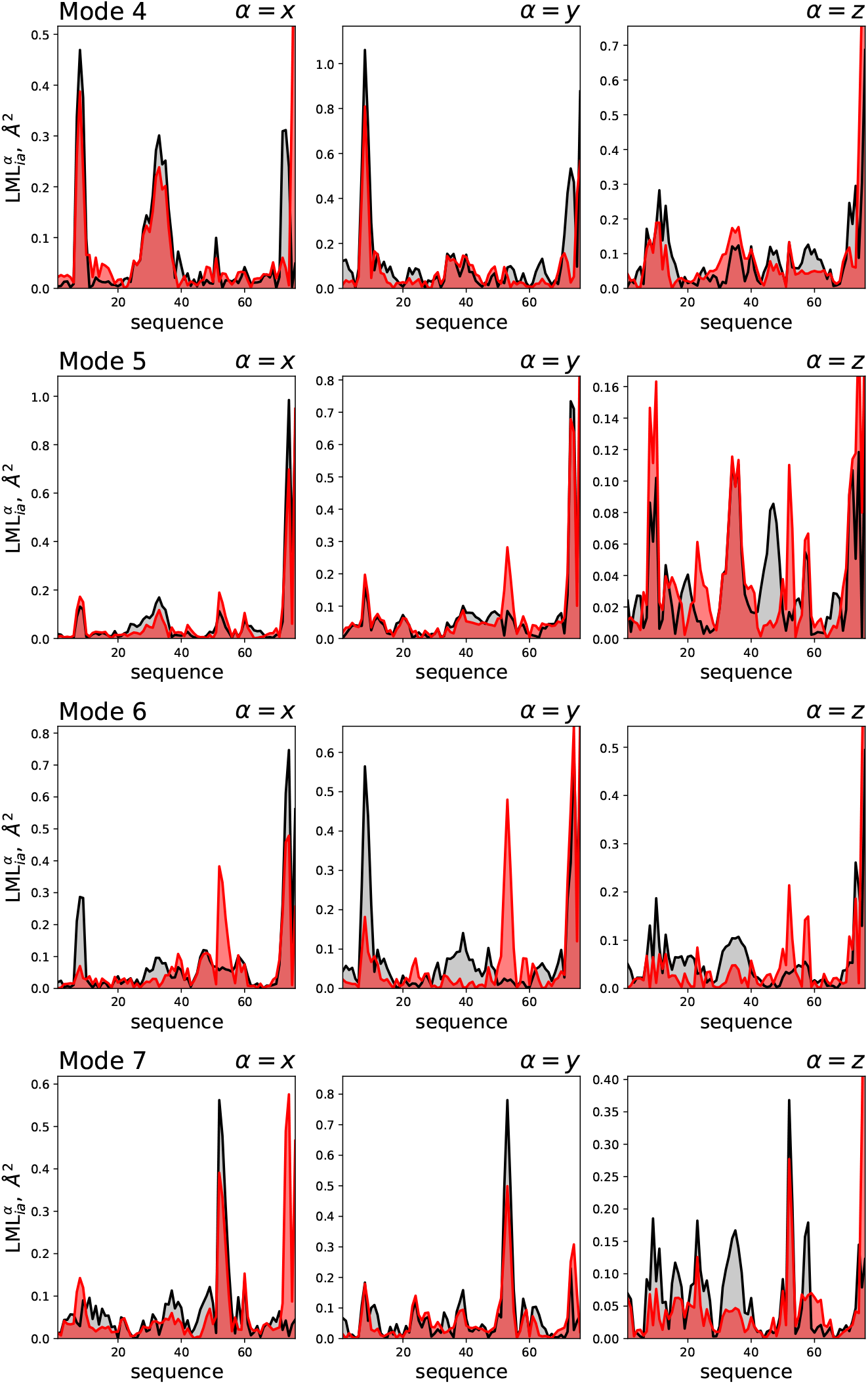
Anisotropic LML, 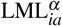 for the next four slowest LE4PD-XYZ modes, as ordered by the *λ_a_* eigenvalues, for the case where hydrodynamic effects are neglected (black) and with hydrodynamic effects included (red).

Note that some modes, such as mode 3, have a FES that is very different if hydrodynamic interaction is included or not. However, the largest fluctuations are localized in the same region of the protein’s primary sequence. Although these modes describe fluctuations in a well-defined regions of ubiquitin (according to Figure 5), the *mechanism* of these dynamics maybe quite different, as indicated by the change in the free-energy landscape.

The situation is different for mode 5, where the anisotropic LML shows differences in the localization of the fluctuations when the two theories are compared. When hydrodynamics are included, there are larger y-coordinate fluctuations predicted in the 50 s loop of ubiquitin not seen when hydrodynamics are neglected. This situation is seen as well in mode 6, although there including hydrodynamics also increase the amplitude of the fluctuations in the C-terminal tail in the y- and z-coordinates, and reduces fluctuations of the Lys11 loop region in the x- and y-coordinates.

Additional changes in the conformational fluctuations described by the two approaches are observed when examining the transition pathways between minima on the free-energy surfaces when hydrodynamic effects are either included or neglected. Figure 11 demonstrates how hydrodynamic effects can alter the predicted conformational changes along mode-dependent transition pathways. For the slowest LE4PD-XYZ mode, shown in Figure 11a, there is little change in the free-energy surface when hydrodynamic effects are included and hence little change in either the predicted transition pathway between minima or the corresponding conformational fluctuations undergone by the protein.

**FIG. 11.**
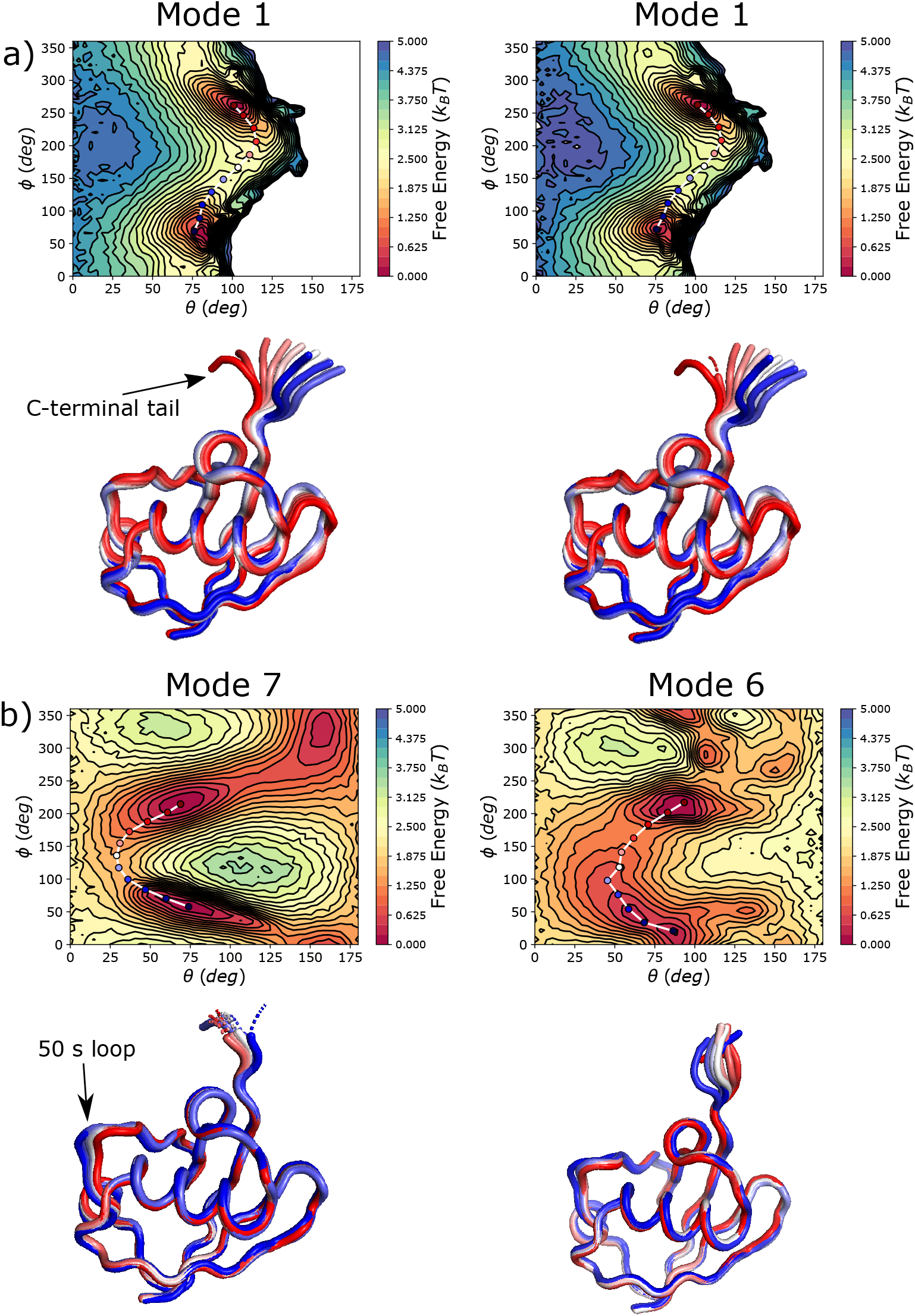
Comparing the mode-dependent fluctuations along a path in the free-energy surface for the slow LE4PD-XYZ modes when hydrodynamics are neglected (left), which is equivalent to PCA, or included (right) for a) LE4PD-XYZ mode 1 and b) LE4PD-XYZ mode 7 without HI (left) and mode 6 with HI (right). Representative structures of ubiquitin for each image along the pathway are given below the corresponding free-energy surface, with the colors of the structure identical to the corresponding image along the pathway.

However, for higher order modes, where the eigenvectors show disagreement between the two approaches, the free-energy landscape is strongly modified and we observe corresponding alterations of the conformational pathways crossing the barriers between energy minima, and related changes in conformational fluctuations along those pathways. Figure 11b shows this effect for mode 7 without hydrodynamics and mode 6 with hydrodynamics (which is the mode with the highest overlap with mode 7 without hydrodynamics). While the free-energy surfaces look roughly the same, there are significant differences in terms of barriers along the pathway between minima and the number of ‘trap states’ along the pathway. Furthermore, mode 7 without hydrodynamics predicts large-scale fluctuations in the C-terminal tail, the 50 s loop, and the Lys11 loop of ubiquitin, while mode 6 with hydrodynamics only predicts motion in the C-terminal tail and lower magnitude fluctuations in the 50 s loop. The motion of the C-terminal tail is quite different in the two modes.

## VII. KINETICS OF BARRIER CROSSING IN THE MODE-DEPENDENT FREE ENERGY LANDSCAPE, CALCULATED BY MARKOV STATE MODELS: A COMPARISON OF PCA VERSUS THE DIFFUSIVE LANGEVIN APPROACH OF THE LE4PD-XYZ

The examples illustrated in the previous section show how examining the fluctuations predicted from the two different approaches provides important insights on the relevance of hydrodynamic interaction in the mode decomposition of the protein dynamics. Here we evaluate the quantitative timescale of the protein fluctuations by analyzing the dynamics of barrier crossing with a Markov State Model analysis.

Markov state models (MSMs) are discrete-state master equation used to determine the kinetics of processes in multi-dimensional energy landscapes.^54^ It is convenient to simplify the analysis of the free energy surface by mapping the multi-dimensional free energy landscape into a set of slow variables in the state space. Those slow variables are extracted from a time-ordered set of configurations, usually generated by an MD simulation, using a procedure of dimensionality reduction like PCA^26,54,55^. In the MSM approach, the state space of slow variables from the simulation is broken into a set of *L* discrete states. The conditional probability of transitioning between two of the given states *i* and *j* is calculated from the MD trajectory by sampling it at a properly-selected lagtime, *τ*. The transition probability is stored in a transition matrix, **T** (*τ*), as *T_ij_*(*τ*). The transition matrix is diagonalized to obtain a set of eigenvalues and eigenvectors; because **T** (*τ*) is a stochastic matrix, its eigenvalues are bounded from above by 1 and all other eigenvalues are of modulus strictly less than 1, according to Perron’s theorem.56 Following the same theorem, the first right eigenvector, *ψ*_1_, of the transition matrix is a vector of “1’s”, *ψ*_1_ = (1, 1,…, 1)^*T*^ and the first left eigen-vector, *ϕ*_1_, is equal to the stationary distribution of the system, *π*; *ϕ*_1_ = *π*.^46,56^

Using the relationship of **T** (*τ*) to the corresponding rate matrix, **K** (*τ*), **T** (*τ*) = *e*^**K**(*τ*)*τ*^, the eigenvalues 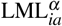 of the transition matrix can be used to find the timescales, *t_i_*, of the dynamic processes described by the MSM^26,54,55^:

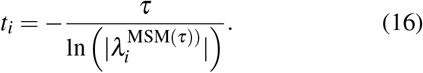

This definition of the timescale of the transition relies on the Chapman-Kolmogorov theorem of Markovian statistics.^54^ Since the eigenvalues of **T** (*τ*) are sorted in descending order, 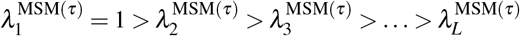, and 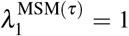 corresponds to the stationary distribution. The slowest non-stationary process described by the MSM corresponds to *ψ*_2_ with a timescale 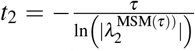. Thus, the slowest MSM mode, *t*_2_, will give the timescale of barrier crossing in the free energy landscape. However, as aforementioned, the MSM analysis is more conveniently applied when the multidimensional free energy landscape is reduced into three dimensional free energy maps. The dimensionality reduction is often performed by using PCA normal modes. However, this study has shown that adopting an anisotropic LE4PD normal mode analysis can give fluctuations that properly include the effect of the solvent and, thus, are more physically sound.

The LE4PD formalism has the advantage of decomposing the complex dynamics of a protein in modes that are quasi-linearly-independent. They provide free energy landscapes that are dimensionally reduced and whose linear combination reconstructs the complete dynamics of the protein.

However, the application of the MSM requires the presence of high enough energy barriers, so that it is possible to separate fast interconverting motions that occur in the energy wells from the slow transitions due to crossing of barriers between states. We have observed that while the slowest protein modes have high energy barriers, and the most local modes have those as well, the intermediate LE4PD modes present a less structured energy landscape, with not localized high energy barriers.^18^ In this study, the MSM analysis is applicable to the ten slowest dynamical modes. Higher index modes show a rough free energy landscape, which is well approximated by a diffusive approach to barrier crossing, such as Kramers equation.^47^

For the slowest LE4PD-XYZ modes, the effective kinetics is determined by performing the MSM analysis in each mode’s FES in (*θ_a_*(*t*)*, ϕ_a_*(*t*)) coordinate space. The second MSM eigenvector, *ψ*_2_, separates the FES into macrostates where the protein rapidly interconverts, while transitions between macrostates are slow. The trajectories are sampled at a lag time, *τ*, that is consistent with the Markovian statistics, as tested using the Chapman-Kolmogorov criteria. In particular, we adopted a method that use the committor function to identify the top of the transition barrier and that we proposed in a recent publication.^18^ In our method the lagtime *τ* in the MSM are selected based on the projection of *ψ*_2_ onto the (*θ_a_, φ_a_*) surface. The longest lagtime *τ* for which the maximum and minimum projections of *ψ*_2_ were both located in deep minima on the surface were chosen as the lagtimes for the MSM on the slow LE4PD-XYZ modes, both with and without HI. This method of selecting *τ* effectively places the node of *ψ*_2_ at the top of the largest barrier on the (*θ_a_, ϕ_a_*) surface. So, the *t*_2_ of the MSM corresponds to the timescale it takes the system to move from one minimum on the surface to the other over the barrier. If the kinetics on the surface follow two-state kinetics, then the locations where *ψ*_2_ has a node, discrete states where *ψ*_2_ = 0, and there the committor will equal 0.5,^27,28^ which is the value for a transition state on the surface.^27,57^ In that case, *ψ*_2_ and the committor give the same information.

Furthermore, for all the MSMs presented here, the number of discrete states *L* = 1000, which is selected based on crossvalidation analyses^58–60^ performed on the presented MSMs. **T**(*τ*) is constructed using the reversible, maximum likelihood estimator given in^61^. An in-depth example of how the MSM is constructed for LE4PD-XYZ mode 7 is given in the Supplementary Material.

Table I shows the timescales for the slowest-occurring processes in the (*θ_a_*(*t*)*, ϕ_a_*(*t*))-space for the ten slowest LE4PD-XYZ modes, when hydrodynamic effects are included and when the HI contributions are neglected. For the LE4PD-XYZ with HI, the two slowest modes, as predicted by the MSM, are modes 1 and 4, which correspond to high-amplitude motions in the the C-terminal tail and the Lys11 loop of ubiquitin occurring over a timescale of 8.0 and 6.4 ns, respectively.

**TABLE 1.**
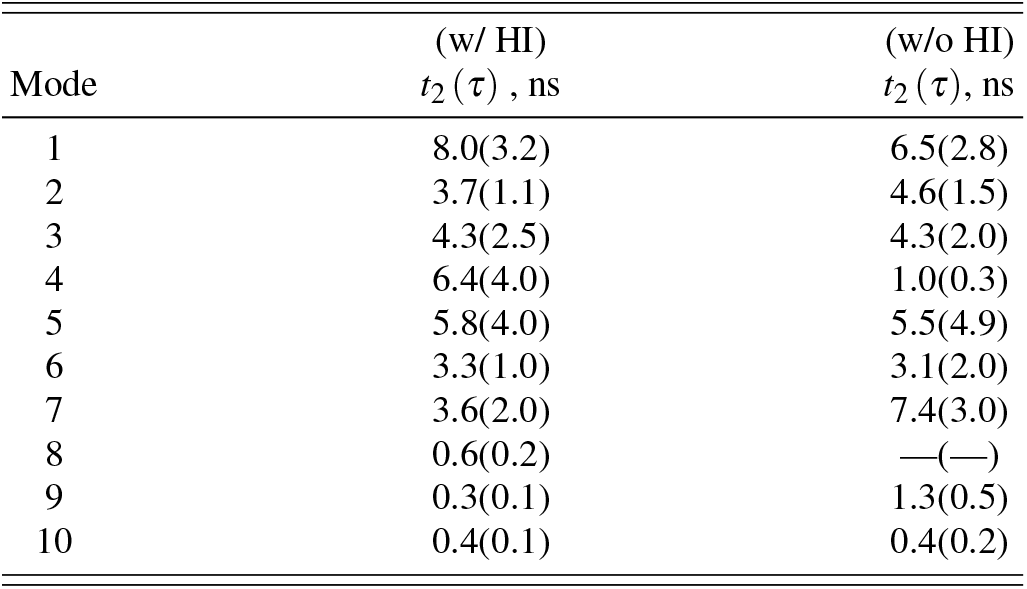
Lagtimes, *τ*, and predicted timescales of the slowest process from the MSM, *t*_2_, (both in ns) of the ten slowest LE4PD-XYZ with and without hydrodynamic interaction included.

Table I also shows the lagtimes and the slowest timescales from a MSM constructed on the two-dimensional FES of the ten slowest LE4PD-XYZ modes without HI, which would correspond to PCA. Qualitatively, the timescales for the slow modes without HI are similar to those with HI. One exception is mode 4, which is predicted to occur on a much faster timescale when HI are neglected, likely due to the lack of motion in the C-terminal tail compared with the same index mode when HI is included. Also, mode 8 without HI occurs on a timescale that is too fast to be modeled with an MSM. Finally, without HI, the LE4PD-XYZ method predicts the slowest mode is mode 7 with a timescale of 7.4 ns. Mode 7 describes motion almost exclusively in the 50 s loop of ubiquitin. Although slightly faster than the 10 ns timescale predicted for a similar dynamics in the isotropic LE4PD theory,^18^ nevertheless the qualitative result is the same, in that there is a single, slow mode that isolates the slowest dynamics in the 50 s loop of ubiquitin. The LE4PD-XYZ theory predicts that this characteristic motion of the 50 s loop is split between modes 6 and 7, which is why the approach with HI does not select this motion as the single slowest mode (Figure 10).

Thus, it is interesting to notice how the inclusion of the hydrodynamic interaction modifies both the energy maps and the timescales of fluctuations as measured by Markov State Model analysis. Furthermore, the slow modes identified by the anisotropic LE4PD and PCA display a dynamics in the 50 s loop of ubiquitin, which is in agreement with the results of the isotropic LE4PD equation,^18^ while the LE4PD-XYZ with hydrodynamics identifies as the slowest fluctuation the dynamics in the C-terminal tail of ubiquitin, which is the second slowest motion, as identified by the isotropic LE4PD.^18^

## VIII. COMPARING THE TIMESCALES PREDICTED BY THE DECAY OF THE MODE TIME CORRELATION FUNCTION

A common method used to calculate the timescales for the decay of the PCA modes is the integration of the time correlation function for each mode,^10,19^ defined as:

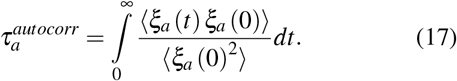

This approach gives the decorrelation time for any arbitrary stochastic process.^62^ In practicality, the upper limit of the integral is taken to be the lagtime *t* where the autocorrelation function hits 0 for the first time.^10,19^ If the process is characterized by a single exponential decay, then

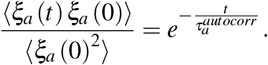

However, in general, for PCA (or LE4PD) modes calculated from a long equilibrium MD simulation of a folded protein, such as the 1-microsecond simulation analyzed here, the relaxation spectrum of the mode autocorrelation function will be more complicated than single exponential.^63,64^ Thus, 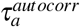 will give an averaged value of the timescale, which includes many relaxation processes.

Using the slowest timescale from the MSM constructed on the (*θ_a_*, *φ_a_*) surfaces also estimates the slowest timescale process of each *ξ_a_*(*t*), but makes the assumption that the kinetic process is Markovian. In general, we do not expect for the two timescales to be identical.

Here, the timescale 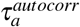 is compared to the previously estimated mode-dependent timescales, namely the *t*_2_ from the MSM (along with the MSM lagtime, *τ*^MSM^, used to generate the associated MSM) and the diffusive *τ_a_* timescale from the LE4PD-XYZ equation of motion, for the ten slowest LE4PD-XYZ modes, either without (Table II) or with (Table III) hydrodynamic interactions included. For all the modes shown here, the *τ_a_* calculated from the equation of motion 6 are lower bounds to *t*_2_ and 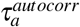, as expected since the *τ_a_* do not account for free-energy barriers along the mode coordinate. In general, for the slowest modes, the timescales calculated using either the MSM or the autocorrelation function are in reasonable agreement, especially for modes 1, 3, and 5 without HI and modes 1, 4, and 5 with HI.

**TABLE 2.** Timescales for the first ten LE4PD-XYZ mode without hydrodynamics. Barrier-free timescales predicted from the LE4PD-XYZ equation, *τ_a_*; the slowest process of the Markov state model, *t*_2_ (with the lagtime of the Markov state model, *τ*, given in parentheses next to *t*_2_); and the de-correlation timescale, 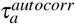, from intergrating the normalized autocorrelation function of the LE4PD-XYZ modes. All timescales are in ns.

| Mode | $\tau_a$, ns | $t_2$ ($\tau$), ns | $\tau_a^{autocorr}$, ns |
| --- | --- | --- | --- |
| 1 | 2.51 | 6.5(2.8) | 9.2 |
| 2 | 1.17 | 4.6(1.5) | 10.2 |
| 3 | 0.52 | 4.3(2.0) | 9.0 |
| 4 | 0.24 | 1.0(0.3) | 5.5 |
| 5 | 0.16 | 5.5(4.9) | 8.5 |
| 6 | 0.15 | 3.1(2.0) | 4.2 |
| 7 | 0.11 | 7.4(3.0) | 44.9 |
| 8 | 0.09 | —(—) | 2.0 |
| 9 | 0.08 | 1.3(0.5) | 5.5 |
| 10 | 0.06 | 0.4(0.2) | 7.1 |

**TABLE 3.** Timescales for the first ten LE4PD-XYZ mode with hydrodynamics. Barrier-free timescales predicted from the LE4PD-XYZ equation, *τ_a_*; the slowest process of the Markov state model, *t*_2_ (with the lagtime of the Markov state model, *τ*, given in parentheses next to *t*_2_); and the de-correlation timescale, 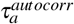, from inter-grating the normalized autocorrelation function of the LE4PD-XYZ modes. All timescales are in ns.

| Mode | $\tau_a$, ns | $t_2$ ($\tau$), ns | $\tau_a^{autocorr}$, ns |
| --- | --- | --- | --- |
| 1 | 0.92 | 8.0(3.2) | 9.8 |
| 2 | 0.46 | 3.7(1.1) | 13.8 |
| 3 | 0.20 | 4.3(2.5) | 10.7 |
| 4 | 0.15 | 6.4(4.0) | 6.0 |
| 5 | 0.11 | 5.8(4.0) | 11.7 |
| 6 | 0.10 | 3.3(1.0) | 31.1 |
| 7 | 0.096 | 3.6(2.0) | 15.7 |
| 8 | 0.057 | 0.6(0.2) | 3.1 |
| 9 | 0.054 | 0.3(0.1) | 2.0 |
| 10 | 0.051 | 0.4(0.1) | 8.4 |

The main discrepancies arise for modes 8 through 10 in both cases, which are the modes where the MSM starts to become less effective and the dynamics approach a regime where the crossing of energy barriers in the (*θ_a_*, *φ_a_*) surfaces becomes more diffusive due to the rough free-energy landscape.^18^ For mode 7 without HI and modes 6 and 7 with HI, which all describe the slow motion in the 50 s loop of ubiquitin, the autocorrelation function relaxes more slowly than the timescale predicted by the MSM. In this case, the difference is likely due to the methodology used to parameterize the MSMs, where the lagtime of the MSM *τ* is selected such that the slowest timescale of the MSM describes transitions between the minima on the surface. In fact the mode trajectory samples not only transitions between the minima but also rare dynamics in the highest energy regions of the surface. These rare events may occur over even longer timescales. Since the autocorrelation function of *ξ_a_* accounts for *all* the processes occurring, 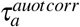 inherits this information and reports longer timescales than the MSM, in general.

## IX. DISCUSSION AND CONCLUSIONS

Large-scale, anisotropic fluctuations in protein dynamics are important as they lead to rare conformational transitions, which are deemed to be relevant for protein folding, and, more generally, for the protein’s biological function.^65,66^ A popular method to select these large fluctuations is a Principal Component Analysis or PCA.^9^ In an MD trajectory, PCA identifies collective fluctuations, which are ordered by their decreasing amplitude, from the most extended to the smallest amplitude. While PCA is both computationally convenient and conceptually simple, it lacks a physical basis beyond the empirical observations that it describes some large-scale, collective motions, functional for the protein.^7,11,12^

In this study, we revisit the PCA formalism and formally connect it to a Langevin equation of motion, which was developed to identify slow dynamical modes and study their kinetics in protein dynamics, called the Langevin equation for protein dynamics, or LE4PD.^13–18^ Like the PCA, the LE4PD decomposes the protein’s motion into an orthogonal set of collective coordinates or modes.

To make a formal connection with PCA, the original LE4PD was extended in this study to describe the anisotropic fluctuations around an average structure. We call this formalism the LE4PD-XYZ. This equation of motion, which is solved analytically into eigenvalues and eigenvectors, captures the anisotropic slow fluctuations of a protein’s alpha carbons, starting from the analysis of the atomistic MD trajectory. The LE4PD-XYZ is a first-principles approach, which allows us to formally connect fluctuations to the different force contributions that model proteins’ dynamics. In this way, the LE4PD-XYZ can be viewed as a powerful equation of motion to accurately describe the dynamics of proteins in solutions.

The LE4PD-XYZ is a coarse-grained approach to protein dynamics and describes the slow fluctuations of the alpha-carbons’ coordinates for each residue. All the residues are modeled as interacting via a harmonic potential of mean force, which is built using the covariances of each residue, as calculated from an MD simulation. This anisotropic equation of motion for the fluctuations of the amino acids’ positions (Eq.1) is diagonalized into a set of equations describing the independent, uncorrelated LE4PD-XYZ normal modes (Eq. 6).

This study shows that the anisotropic Langevin equation for protein dynamics, or LE4PD-XYZ, becomes formally equivalent to an equation of motion guided by the forces in the covariance matrix. Thus the LE4PD-XYZ is equivalent to a Principal Component Analysis approach, but only when two specific approximations are adopted. The first approximation is that the equation-of-motion disregards the hydrodynamic interaction, i.e. that there is no correlation in the dynamics of the amino acids caused by the presence of long-ranged interactions mediated by the solvent. This is the so-called “free-draining” limit. The second approximation is that every amino acid in the protein has identical friction. Only when these two approximations are enforced, the fluctuations identified by the PCA become identical to the ones modeled by a diffusive equation of motion. Unfortunately, these approximations are in general not justified, even if they are frequently adopted. Including solvent-mediated interactions is important when one models protein dynamics in an effective medium. And this study shows, specifically, that hydrodynamic interactions modify the dynamics of the protein, and importantly the timescales of the slow modes. Likewise, the degree of exposure to the solvent of each amino acid, and their unique friction, affects the timescale of the amino acids’ dynamics and their fluctuations.

While the dynamics of protein is anharmonic, the Langevin formalism and PCA both rely on the harmonicity of the fluctuations, which is a valid approximation only in the proximity of the folded state. This approximation is common in many structural approaches such as the Gaussian Normal Modes method.^67–69^ However, in the LE4PD approach the anharmonic fluctuations are identified through a procedure that maps the simulation trajectory onto its mode-dependent two-dimensional free energy landscapes. This convenient procedure of variable reductions allows one to study the dynamics of the protein in separated modes. For each mode, one can study the position, amplitude, and timescale of the fluctuations that are important for the biological function.^18^

The eigenvalues of the LE4PD-XYZ define an ‘original’ timescale, which is corrected by the analysis of the mode-dependent FES to include the slowing down of the dynamics due to the presence of high energy barriers. The corrected timescale is then calculated either from a Markov State Model analysis of the FES, or from the integral of the time-correlation function of the modes, which is a procedure often used in the PCA.^19^

The study applies the LE4PD-XYZ method to a 1-*μ*s simulation of ubiquitin and compares the results with an analysis performed using the Principal Component Analysis method. The comparison between the timescale from the eigenvalues, and the more realistic timescales measured by applying a Markov state model analysis to the mode-dependent free energy maps, after identifying the leading transition pathways for these fluctuations, show the relevance of the energy barriers in measuring the kinetic timescale of fluctuations (see for example Tables I, II, and III).

Interestingly, the decay of the time-correlation function of the mode coordinates, which is the procedure often used to calculate the timescale in PCA, is qualitatively consistent the more elaborate Markov state model analysis of the slow pathways in the LE4PD-XYZ free energy maps. This result confirms the need to include both barrier crossing and the hydrodynamic interaction in a Langevin description of protein dynamics.

Furthermore, when we examined the effect including hydrodynamics on the predicted mode-decomposition of the dynamics, we observed that some, but not all, of the slowest modes are little changed. However, the introduction of hydrodynamic interaction has important effects on the faster modes, which are involved in local-scale processes of the proteins, for example in chemical reactions.

Specifically for ubiquitin, we observe that the slowest LE4PD-XYZ modes predict timescales between 300 ps and 8 ns, roughly in-line with those predicted in an early study of the same system using the isotropic LE4PD.18 In contrast, the anisotropic LE4PD-XYZ approach is not able to isolate the slow fluctuations of the 50 s loop of ubiquitin, seen in mode 9 by the isotropic approach. The anisotropic description separates into the directional contributions this slow dynamics. However, when the HI is neglected one can again detect the unidirectional, slow fluctuation of the 50 s loop.

For ubiquitin, both the LE4PD-XYZ and PCA identify slow fluctuations and large-amplitude motion in the C-terminal tail region. It is known that the tail of ubiquitin is involved in many of the protein’s binding events to substrates, both covalent^70,71^ and non-covalent. ^72^ The wide range of possible conformations that are available for the binding of the tail may be important for the protein to discriminate among different possible reaction substrates. Thus, in line with the conformational selection hypothesis,^73,74^ the large number of modes dedicated to describing motion in the C-terminal tail may indicate the opportunity for the protein to follow different transition pathways in the mechanism of binding to different substrates.

In conclusion, we have presented here the formalism for an anisotropic Langevin equation, the LE4PD-XYZ, that describes the motions of a protein in terms of a set of orthogonal normal modes and used it to analyze a 1-*μ*s MD simulation of the protein ubiquitin. This approach coarse-grains the dynamics of the protein at the level of the protein’s amino acids and accounts for the hydrodynamic interaction (HI) between amino acids as well as free-energy barriers along each of the LE4PD-XYZ modes. When HI, the specificity of each amino acid’s friction coefficient, and free-energy barriers are neglected, the LE4PD-XYZ approach maps *exactly* onto the analogous PCA (in the sense that the dynamics is described by the same set of eigenvalues and eigenvectors). The inability of PCA alone to describe the dynamics correctly (unless hydrodynamics and energy barriers are included in the related equation of motion) is highlighted in Figure 1, where the time correlation function calculated from the simulation is compared with the mode-dependent decay of the PCA eigenvalues and displays a clear disagreement, with the correlation functions predicted from the PCA modes decaying too quickly relative to the correlation functions calculated from the simulation trajectory.

This study shows that the inclusion of the HI modulates the location and amplitude of the predicted fluctuations (Figures 5, 9, 10), eigenvalues (Figure 4), free-energy surfaces (Figure 7, 8, 11), and timescales (Table I). Finally, we have also shown how including free-energy barriers causes the dynamics predicted by the slow LE4PD-XYZ modes without HI to be different from those predicted by the analogous PCA (Figures 2, 3). These results demonstrate the importance of considering both hydrodynamic effects, with specific friction coefficients, and energetic barriers to transport when analyzing the equilibrium dynamics of ubiquitin about its folded state. Only by including these effects the time correlation functions that define the decay of local correlations quantitatively reproduce the decay measured in the atomistic simulations.

## SUPPLEMENTARY MATERIAL

The Supplementary Material contains a detailed derivation of the **H** for the LE4PD-XYZ, a formal relationship between the **U** matrices from the isotropic LE4PD and the LE4PD-XYZ, comparisons of the conformational fluctuations predicted along the interpolated and minimum energy pathways for more of the slow LE4PD-XYZ modes, and a detailed example of the MSM construction for the first LE4PD-XYZ mode.

## DEDICATION

M.G.G. dedicates this paper to all the women (friends, family, and colleagues) who made her life and her work stronger and more enjoyable with their advice, love, and support.

## ACKNOWLEDGMENTS AND FUNDING

E.R.B. was supported by the John Keana Graduate Student Fellowship from the University of Oregon and the National Science Foundation through grants CHE-1665466 and CHE-1362500 to M.G.G. The computational work was performed on the supercomputer Comet at the San Diego Supercomputer Center, with the support of XSEDE^75^ allocation TG-CHE100082 to M.G.G. (XSEDE is a program supported by the National Science Foundation under Grant No. ACI-1548562).

## DATA AVAILABILITY

The codes used to perform the LE4PD-XYZ analysis described here are available on GitHub (https://github.com/GuenzaLab/LE4PD-XYZ), and the processed MD trajectory and 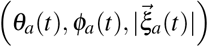trajectories for the first 10 LE4PD-XYZ modes both with and without hydrodynamics included are available on Zenodo (https://doi.org/10.5281/zenodo.4312224).

