## Supplementary Materials for "Comparison between slow, anisotropic LE4PD fluctuations and the Principal Component Analysis modes of Ubiquitin"

(Dated: December 21, 2020)

*.

### I. ANISOTROPIC HYDRODYNAMICS

In this section, we propose an extension of the traditional derivation of the preaveraged hydrodynamic interaction while including the anisotropic formalism. The derivation is straightforward, but it may be useful, as we are not aware that it has been reported previously.

The system we are modeling is a coarse-grained description of a protein in solution, where the protein is a chain of beads, or friction points, in an effective solvent. Each bead represents one amino acid along the protein's primary sequence. The equation of motion of the fluctuations for a protein consisting of N beads, while neglecting hydrodynamic effects, is

$$\bar{\zeta}\Delta\dot{R}_i^{\alpha}(t) = -k_B T \sum_{\beta\in\{x,y,z\}} \sum_{j=1}^{N} A_{ij}^{\alpha\beta}\Delta R_j^{\beta}(t) + F_i^{\alpha}(t), \tag{S1}$$

where $\bar{\zeta}$ is the average bead friction coefficient, $\bar{\zeta} = \frac{1}{N}\sum_{i=1}^{N}\zeta_i$, $\Delta R(t)$ is the 3N x 1 column vector of Cartesian displacements of each bead from its equilibrium position:

$$\Delta R(t) = \left[x_1(t) - \langle x_1\rangle, y_1(t) - \langle y_1\rangle, z_1(t) - \langle z_1\rangle, x_2(t) - \langle x_2\rangle, \ldots, z_N(t) - \langle z_N\rangle\right]^T, \tag{S2}$$

with $\Delta\dot{R}_i^{\alpha}(t)$ denoting the fluctuation of the $\alpha$-component of alpha-carbon $i$ away from its equilibrium position; $k_B$ is Boltzmann's constant, $T$ is the temperature, $A$ is a 3N x 3N structural matrix defining the coupling between the fluctuations of each component of each alpha-carbon from its average position, and $F_i^{\alpha}(t)$ is a stochastic force along the $\alpha$ component that represents the fast collisions between the solvent and the $i^{th}$ bead that obeys a delta-correlated, white-noise fluctuation-dissipation theorem:

$$\begin{aligned} \langle F_i^{\alpha}(t)\rangle &= 0 \\ \langle F_i{}^{\alpha}(t)F_j{}^{\beta}(t')\rangle &= 2k_B T\bar{\zeta}\delta(t-t')\delta_{ij}\delta_{\alpha\beta}, \end{aligned} \tag{S3}$$

with $\delta_{ij}, \delta_{\alpha\beta}$ Kronecker delta symbols and $\alpha, \beta$ labeling the Cartesian indices of the force, $\alpha, \beta \in \{x, y, z\}$.

Since the protein is surrounded by a solvent, accounting for hydrodynamic effects is crucial for an accurate prediction of the protein's dynamic and kinetic behavior. The presence of the protein in the solvent will perturb the solvent's velocity; treating each bead in the protein as a solid sphere of radius $s_i$, the velocity of the $\alpha$ component of the fluid at the location of bead $i$ is given by

$$v_i^{\alpha} = v_i^{\alpha,0} + v_i^{'\alpha}, \tag{S4}$$

where $v_i^{\alpha,0}$ is the unperturbed $\alpha$ component of the velocity of the solvent (the velocity it would possess in the absence of the protein) and the perturbation in the velocity due to the presence of the other $j \neq i$ beads in the protein:

$$v_i^{'\alpha} = \sum_{\beta\in\{x,y,z\}} \sum_{j} T_{ij}^{\alpha\beta} F_j{}^{\beta,\zeta}. \tag{S5}$$

As with $\Delta R(t)$, $v'$ is an 3N x 1 column vector, with $v_i^{'\alpha}$ describing the perturbed velocity at alpha-carbon $i$ in along the $\alpha$ component. $T_{ij}^{\alpha\beta}$ is an element of a $3N \times 3N$ tensor $\mathbf{T}$ that describes the coupling between the force exerted by bead $j$ along the $\alpha$ component of the solvent and the resulting velocity perturbation experienced by bead $i$ along the $\beta$ component, and $F_j{}^{\alpha,\zeta}$ is the total force exerted on the solvent along component $\alpha$ by bead $j$. In equation S5, the exact solution for $T_{ij}^{\alpha\beta}$ is (see, e.g. [1]),

$$\begin{aligned} T_{ij}^{\alpha\beta} &= \frac{1}{8\pi\eta_w r_{ij}}\left(\delta_{\alpha\beta} + (\widehat{r}\widehat{r})_{\alpha\beta} + \frac{s_i^2 + s_j^2}{r_{ij}^2}\left[\frac{1}{3}\delta_{\alpha\beta} - (\widehat{r}\widehat{r})_{\alpha\beta}\right]\right), \ i \neq j \\ T_{ij}^{\alpha\beta} &= 0, \ i = j. \end{aligned} \tag{S6}$$

In equation S6, $r_{ij}$ is the distance between beads $i$ and $j$, $s_i$ is the radius of bead $i$, and $\widehat{r}$ is a unit vector in the direction of $r_{ij}$, $\widehat{r} = \frac{\mathbf{r}}{r_{ij}} = \left(\frac{x_{ij}}{r_{ij}}, \frac{y_{ij}}{r_{ij}}, \frac{z_{ij}}{r_{ij}}\right)^T$. Accounting for the perturbation in the velocity at the location of bead $i$ and the individual friction coefficient of each bead gives a modified Langevin equation:

$$\zeta_i\left(\Delta\dot{R}_i^{\alpha}(t) - v_i{}^{0,\alpha} - v_i^{'\alpha}\right) = -k_B T\sum_{\beta}\sum_{k} A_{ik}^{\alpha\beta}\Delta R_k^{\beta}(t) + F_i^{\alpha}(t)$$

$$\zeta_i \left( \Delta \dot{R}_i^{\alpha}(t) - \sum_{\beta} \sum_{j} T_{ij}^{\alpha\beta} F_j^{\beta,\zeta} \right) = -k_B T \sum_{\beta} \sum_{k} A_{ik}^{\alpha\beta} \Delta R_k^{\beta}(t) + F_i^{\alpha}(t). \tag{S7}$$

In the second line, it has been assumed that no external force has been applied to the solvent (e.g. there are no plates at the top and bottom of the box shearing the fluid) so that $v_i^0 = 0$ and, equation S5 has been used to write $v_i'$ in terms of the forces exerted on the solvent by the beads. In the bead-and-spring-based model presented here, the beads can exert only two forces on the solvent: a spring force defined by $A$, $F_{\text{Spring},ij}^{\beta}(t) = -k_B T \sum_{\gamma} \sum_j A_{ij}^{\beta\gamma} \Delta R_j^{\gamma}(t)$, and a stochastic force due to random, thermal fluctuations of the solvent moving the beads, $F_j(t)$. Thus,

$$\begin{aligned} F_i^{\beta,\zeta} &= \sum_j F_{\text{Spring},ij}^{\beta} + F_i^{\beta}(t) \\ &= -k_B T \sum_{\gamma} \sum_{j} A_{ij}^{\beta\gamma} \Delta R_j^{\gamma}(t) + F_i^{\beta}(t). \end{aligned} \tag{S8}$$

Substituting equation S8 into equation S7 gives the explicit (anisotropic) equation of motion for bead $i$:

$$\zeta_i \left( \Delta \dot{R}_i^{\alpha}(t) - \sum_{\beta} \sum_{j} T_{ij}^{\alpha\beta} \left[ \sum_{\gamma} \sum_{k} -k_B T A_{jk}^{\beta\gamma} \Delta R_k^{\gamma}(t) + F_j^{\beta} \right] \right) = -k_B T \sum_{\beta} \sum_{k} A_{ik}^{\alpha\beta} \Delta R_k^{\beta}(t) + F_i^{\alpha}(t). \tag{S9}$$

To formulate a hydrodynamic interaction matrix, $H$, that describes the effect that hydrodynamics has on the motions of the other beads in the protein, there are two cases of equation S9 to treat:

1. $i = j$: In this situation, $T_{ij}^{\alpha\beta} = 0$, so equation S9 reduces to

$$\zeta_i \Delta \dot{R}_i^{\alpha}(t) = -k_B T \sum_{\beta} A_{ik}^{\alpha\beta} \Delta R_k^{\beta}(t) + F_i^{\alpha}(t)$$

$$\Rightarrow \Delta \dot{R}_i^{\alpha}(t) = \frac{\bar{\zeta}}{\zeta_i} \left( \frac{-k_B T}{\bar{\zeta}} \sum_{\beta} \sum_{k} A_{ik}^{\alpha\beta} \Delta R_k^{\beta}(t) \right) + \frac{F_i^{\alpha}(t)}{\zeta_i}. \tag{S10}$$

   Equation S10 implies that the form of the original Langevin equation, equation S1, is recovered by defining $H_{ii}^{\alpha\beta}$ as $H_{ii}^{\alpha\beta} = \frac{\bar{\zeta}}{\zeta_i} \delta_{\alpha\beta}$. In equation S10, $\zeta_i$ is the site-specific friction coefficient for residue $i$, which is the sum of two contributions. The first contribution is due to the partial exposure of the amino acid to the solvent, and the second is the friction due to its partial exposure to the hydrophobic core of the protein:

$$\zeta_i = 6\pi \left( \eta_w r_{w,i} + \eta_p r_{p,i} \right), \tag{S11}$$

   where $\eta_w$ is the bulk viscosity of the solvent, $r_{w,i}$ is the effective radius of residue $i$ exposed to the solvent, $\eta_p$ is the viscosity of the protein (the 'internal' viscosity), and $r_{p,i}$ is the effective radius of residue $i$ exposed to the hydrophobic core of the protein. Both $r_{w,i}$ and $r_{p,i}$ are caclulated from the simulation and $\eta_p$ is assumed to be 'slaved' to the solvent viscosity, so that it is proportional to the solvent viscosity. Specifically, it is assumed that the internal viscosity is the solvent viscosity, rescaled by a local-barrier energy scale of $k_B T$: $\eta_p = \exp[k_B T / k_B T] \eta_w \approx 2.71828 \eta_w$.

2. $i \neq j$: In this situation, the force terms on the right-hand side (RHS) of equation S9 have already been treated in the $i = j$ case, so that, when $i \neq j$, the equation of motion reduces to

$$\zeta_i \Delta \dot{R}_i^{\alpha}(t) - \zeta_i \sum_{\beta} \sum_{j} T_{ij}^{\alpha\beta} \left[ \sum_{\gamma} \sum_{k} -k_B T A_{jk}^{\beta\gamma} \Delta R_k^{\gamma}(t) + F_j^{\beta} \right] = 0$$

$$\Rightarrow \Delta \dot{R}_i^{\alpha}(t) = \bar{\zeta} \sum_{\beta} \sum_{j} T_{ij}^{\alpha\beta} \left[ \sum_{\gamma} \sum_{k} -\frac{k_B T}{\bar{\zeta}} A_{jk}^{\beta\gamma} \Delta R_k^{\gamma}(t) + \frac{F_j^{\beta}}{\bar{\zeta}} \right]. \tag{S12}$$

   Again, by inspection, the form of equation S1 is recovered by setting $H_{ij}^{\alpha\beta} = \bar{\zeta} T_{ij}^{\alpha\beta}$.

Combing these two cases, the complete hydrodynamic interaction matrix is defined piecewise as

$$H_{ij}^{\alpha\beta} = \begin{cases} \frac{\overline{\zeta}}{\zeta_i}\delta_{\alpha\beta}, \ i = j \\ \\ \overline{\zeta}T_{ij}^{\alpha\beta}, \ i \neq j \end{cases} \tag{S13}$$

The off-diagonal elements of $H$ can be simplified if we assume that only the portion of each bead exposed to solvent contributes to the hydrodynamic effect; that is, for each $H_{ij}^{\alpha\beta}$, $i \neq j$, we assume $H_{ij}^{\alpha\beta} = \overline{\zeta}_w T_{ij}^{\alpha\beta}$, so that

$$\begin{aligned} H_{ij}^{\alpha\beta} = \overline{\zeta}_w T_{ij}^{\alpha\beta} &= 6\pi\eta_w \overline{r}_w T_{ij}^{\alpha\beta} \\ &= \frac{6\pi\eta_w\overline{r}_w}{8\pi\eta_w r_{ij}}\left(\delta_{\alpha\beta} + (\widehat{r}\widehat{r})_{\alpha\beta} + \frac{s_i^2 + s_j^2}{r_{ij}^2}\left[\frac{1}{3}\delta_{\alpha\beta} - (\widehat{r}\widehat{r})_{\alpha\beta}\right]\right) \\ &= \frac{3\overline{r}_w}{4r_{ij}}\left(\delta_{\alpha\beta} + (\widehat{r}\widehat{r})_{\alpha\beta} + \frac{s_i^2 + s_j^2}{r_{ij}^2}\left[\frac{1}{3}\delta_{\alpha\beta} - (\widehat{r}\widehat{r})_{\alpha\beta}\right]\right) \end{aligned} \tag{S14}$$

When kept in the form given in equation S14, the hydrodynamic interaction matrix is referred to as the *Rotne-Prager tensor*. This form gives the exact solution for the Stokes flow around a solid sphere moving through a viscous, incompressible fluid, and is useful because it accounts for the finite size of the beads. The hydrodynamic interaction matrix can be simplified in the limit of large inter-bead distances, i.e. in the limit of $r_{ij} \to \infty$. In this case, the term within the brackets in equation S14 goes as $\frac{1}{r_{ij}^3}$ and is negligible compared to the first term in equation S14. That is,

$$\begin{aligned} \lim_{r_{ij}\to\infty} H_{ij}^{\alpha\beta} &= \lim_{r_{ij}\to\infty} \frac{3\overline{r}_w}{4r_{ij}}\left(\delta_{\alpha\beta} + (\widehat{r}\widehat{r})_{\alpha\beta} + \frac{s_i^2 + s_j^2}{r_{ij}^2}\left[\frac{1}{3}\delta_{\alpha\beta} - (\widehat{r}\widehat{r})_{\alpha\beta}\right]\right) \\ &= \frac{3\overline{r}_w}{4r_{ij}}\left(\delta_{\alpha\beta} + (\widehat{r}\widehat{r})_{\alpha\beta}\right). \end{aligned} \tag{S15}$$

The hydrodynamic interaction matrix defined in equation S15 is known as the *Oseen tensor* and is simpler than what is given in equation S14. This approximation treats the beads as point particles, and is accurate for large bead separations. However, in the case that the beads approach each other to within a bead radius, i.e. when $r_{ij} \leq \max(s_i, s_j)$, the Oseen tensor can give negative, unphysical eigenvalues because, in that instance, the point particle approximation is not really valid. Regardless if the off-diagonal elements of $H$ are defined using equation S14 or S15, the equation of motion with the inclusion of hydrodynamic effects is

$$\overline{\zeta}\Delta\dot{R}_i^{\alpha}(t) = -k_B T \sum_{\beta,\gamma}\sum_{j,k} H_{ij}^{\alpha\beta} A_{jk}^{\beta\gamma}\Delta R_k^{\gamma}(t) + F_i^{\alpha}(t), \tag{S16}$$

It should be noted that, up to this point, the pre-averaging approximation of Kirkwood and Riseman has not been invoked, so that $H$ is still time-dependent.

#### A. Averaging the Hydrodynamic Interaction

A difficulty with the hydrodynamic interaction matrix given in equation S13 is that it is non-linear in $\Delta R$. Zimm's original solution to this problem was the use of the pre-averaging approximation developed by Kirkwood and Riseman in their treatment of the translational diffusion of polymers. Kirkwood and Riseman's approximation assumes that the inter-bead distribution is Gaussian and that

$$\langle H_{ij}\rangle = \frac{3\overline{r}_w}{4}\left\langle \frac{1}{r_{ij}}\left(\widehat{I} + \widehat{r}_{ij}\widehat{r}_{ij}\right)\right\rangle_{|r_{ij}|,\theta,\phi} = \frac{3\overline{r}_w}{4}\left\langle\frac{1}{r_{ij}}\right\rangle_{|r_{ij}|}\left(\langle\widehat{I}\rangle_{\theta,\phi} + \langle\widehat{r}_{ij}\widehat{r}_{ij}\rangle_{\theta,\phi}\right), \ i \neq j; \tag{S17}$$

the second equality follows because the distribution of $\widehat{r}_{ij}$ is independent of its magnitude, $r_{ij}$; the average is taken over the equilibrium distribution of $\vec{r}_{ij}$, $\Psi(\vec{r}_{ij}) = \left(\frac{3}{2\pi|i-j|l^2}\right)^{\frac{3}{2}} \exp\left[-\frac{3\vec{r}_{ij}^2}{2|i-j|l^2}\right]$, with $l$ the average bond length between beads:

$$\langle \cdots \rangle_{|r_{ij}|,\theta,\phi} = \int_0^{2\pi} d\phi \int_0^{\pi} d\theta \sin\theta \int_0^{\infty} dr_{ij} \cdots \Psi(\vec{r}_{ij})$$
$$= \int_0^{2\pi} d\phi \int_0^{\pi} d\theta \sin\theta \int_0^{\infty} dr_{ij} \cdots \Psi(r_{ij}),$$

where $|r_{ij}| = r_{ij}$. Taking the angular average of $\widehat{r}_{ij}\widehat{r}_{ij}$ yields $\langle \widehat{r}_{ij}\widehat{r}_{ij}\rangle_{\theta,\phi} = \frac{4\pi}{3}\widehat{I}$ due to the tensor identity $\langle \widehat{r}_{ij}^{\alpha}\widehat{r}_{ij}^{\beta}\rangle_{\theta,\phi} = \frac{4\pi}{3}\delta_{\alpha\beta}$, where, as before, $\alpha$ and $\beta$ index the Cartesian components of $\widehat{r}_{ij}$. So,

$$\left(\langle \widehat{I}\rangle_{\theta,\phi} + \langle \widehat{r}_{ij}\widehat{r}_{ij}\rangle_{\theta,\phi}\right) = 4\pi\left(\widehat{I} + \frac{1}{3}\widehat{I}\right) = \frac{16\pi}{3}\widehat{I},$$

and equation S17 simplifies to

$$\langle H_{ij}\rangle = 4\pi\overline{r}_w \left\langle \frac{1}{r_{ij}} \right\rangle_{|r_{ij}|} \widehat{I}$$
$$= \overline{r}_w \left\langle \frac{1}{r_{ij}} \right\rangle \widehat{I} \tag{S18}$$

or

$$\langle H_{ij}^{\alpha\beta}\rangle = \overline{r}_w \left\langle \frac{1}{r_{ij}} \right\rangle \delta_{\alpha\beta} \tag{S19}$$

since $\Psi(r_{ij})$ is independent of the angular integral. Because the 3 x 3 blocks of $H$ on the diagonal are already time-independent (depending only on the individual friction coefficients of each bead), the complete pre-averaged, hydrodynamic interaction matrix between beads $i$ and $j$ is given by

$$\langle H_{ij}\rangle = \frac{\overline{\zeta}}{\zeta_i}\delta_{ij}\widehat{I} + (1-\delta_{ij})\left\langle \frac{1}{r_{ij}} \right\rangle \widehat{I}. \tag{S20}$$

or

$$\langle H_{ij}^{\alpha\beta}\rangle = \frac{\overline{\zeta}}{\zeta_i}\delta_{ij}\delta_{\alpha\beta} + (1-\delta_{ij})\left\langle \frac{1}{r_{ij}} \right\rangle \delta_{\alpha\beta}. \tag{S21}$$

What equation S20 means is that pre-averaging the hydrodynamic interaction, while removing the time-dependence, also removes the anisotropic effects inherent in the hydrodynamic interactions between beads. Although equation S20 was derived using the Oseen tensor in $H$, since $(\langle \widehat{r}_{ij}\widehat{r}_{ij}\rangle)_{\alpha\beta} = \frac{1}{3}\delta_{\alpha\beta}$, a pre-averaging of the Rotne-Prager tensor gives the same result as using the Oseen tensor (because the second term on the RHS of equation S14 contains $\frac{1}{3}\delta_{\alpha\beta} - (\widehat{r}_{ij}\widehat{r}_{ij})_{\alpha\beta}$).

### II. RELATIONSHIP BETWEEN THE ISOTROPIC AND ANISOTROPIC LE4PD

In the isotropic case, the relevant structural matrices are given by using the following definitions of the **U** and **A** matrices:[2, 3]

$$U_{N,ij} = \frac{\left\langle \vec{l}_i \cdot \vec{l}_j \right\rangle}{\left\langle |\vec{l}_i| \right\rangle \left\langle |\vec{l}_j| \right\rangle}$$

$$\mathbf{A}_N = \mathbf{a}^T \mathbf{U}_N^{-1} \mathbf{a},$$

with $\vec{l}_i = \left(r_{x,i}, r_{y,i}, r_{z,i}\right)^T$ the bond vector between beads $i$ and $i+1$. Taking the $ij^{th}$ element of $\mathbf{A}^{-1}$ gives

$$\begin{aligned}
\mathbf{A}_N^{-1} &= \left(\mathbf{M}^T \begin{pmatrix} 0 & 0 \\ 0 & \mathbf{U}_N^{-1} \end{pmatrix} \mathbf{M}\right)^{-1} \\
&= \left(a^T \mathbf{U}_N^{-1} a)\right)^{-1} \\
&= a^{-1} \mathbf{U}_N \left(a^T\right)^{-1} \\
&\Rightarrow A_N^{-1}{}_{ij} \approx l^{-2} \mathbf{a}^{-1} \left\langle \vec{l}_i \cdot \vec{l}_j \right\rangle \mathbf{a}^{T^{-1}} = l^{-2} \left\langle \vec{R}_i \cdot \vec{R}_j \right\rangle \\
&\Rightarrow \operatorname{tr}\left(\mathbf{A}_N^{-1}\right) = l^{-2} \sum_i \left\langle \vec{R}_i \cdot \vec{R}_i \right\rangle = \frac{N}{l^2} \left\langle R_g^2 \right\rangle,
\end{aligned} \tag{S22}$$

with $\vec{R}_i$ the distance vector of bead $i$ from the center-of-mass of the polymer, $\langle R_g^2 \rangle$ the average squared radius of gyration, and $\operatorname{tr}(\mathbf{B})$ the trace of a matrix $\mathbf{B}$. The approximation in the fourth line comes from taking all the average square bond lengths between beads to be equal, which is approximately true due to the stiffness of the peptide bond between the alpha-carbons in a protein. The trace of $\mathbf{A}_N^{-1}$ can be related to the trace of the 3N x 3N $\mathbf{A}$ by noting that $\mathbf{A}$ can be written as the difference of two matrices, since $\mathbf{A}^{-1} = \mathbf{C}$:

$$\begin{aligned}
\mathbf{A}^{-1} &= \mathbf{C} = \left\langle \Delta R \Delta R^T \right\rangle \\
&= \left\langle \left(R - \langle R \rangle\right) \left(R - \langle R \rangle\right)^T \right\rangle \\
&= \left\langle R R^T \right\rangle - \langle R \rangle \langle R \rangle^T \\
&= \mathbf{A}_1^{-1} - \mathbf{A}_2^{-1}.
\end{aligned} \tag{S23}$$

Explicitly,

$$\begin{aligned}
\mathbf{A}_1^{-1} &= \begin{pmatrix} x_1 x_1 & x_1 y_1 & x_1 z_1 & x_1 x_2 & \cdots \\ y_1 x_1 & y_1 y_1 & y_1 z_1 & y_1 x_2 & \cdots \\ z_1 x_1 & z_1 y_1 & z_1 z_1 & z_1 x_2 & \cdots \\ \vdots & \vdots & \vdots & \vdots & \ddots \end{pmatrix} \\
&\Rightarrow \operatorname{tr}\left(\mathbf{A}_1^{-1}\right) = x_1 x_1 + y_1 y_1 + z_1 z_1 + x_2 x_2 + \ldots \\
&= \vec{R}_1 \cdot \vec{R}_1 + \vec{R}_2 \cdot \vec{R}_2 + \ldots \\
&= \sum_i \vec{R}_i \cdot \vec{R}_i = N \langle R_g^2 \rangle \Rightarrow \operatorname{tr}\left(\mathbf{A}_1^{-1}\right) \approx l^2 \operatorname{tr}\left(\mathbf{A}_N^{-1}\right).
\end{aligned} \tag{S24}$$

Or, the trace of $\mathbf{A}_1^{-1}$ is approximately equal to the trace of $\mathbf{A}_N^{-1}$, with the approximation the equality of the bond lengths between each bead. The error in this approximation is about 0.5 %, while the error between $l^2 \operatorname{tr}\left(\mathbf{A}_N^{-1}\right)$ and $\operatorname{tr}\left(\mathbf{A}^{-1}\right)$ is of the order $10^{-3}$%; therefore, the equality holds to within the error of the approximation of equal bond lengths.

The accuracy of the normal modes generated from $\mathbf{A}$ can be established by examining their ability to reproduce the structural properties of the protein chain. For example, the mean-squared radius of gyration can be written in terms of the eigenvectors and eigenvalues of $\mathbf{A_1}$:

$$\begin{aligned}\langle R_g^2 \rangle &= \frac{1}{N}\sum_{i=1}^{N}\left\langle \vec{R}_i \cdot \vec{R}_i \right\rangle \\ &= \frac{1}{N}\sum_{i=1}^{N}\left(\langle R_{i,x}R_{i,x}\rangle + \langle R_{i,y}R_{i,y}\rangle + \langle R_{i,z}R_{i,z}\rangle\right) \\ &= \frac{1}{N}\sum_{i=1}^{N}\left(\langle R_{i,x}R_{i,x}\rangle + \langle R_{i,y}R_{i,y}\rangle + \langle R_{i,z}R_{i,z}\rangle\right) \\ &= \mathrm{tr}\mathbf{A_1^{-1}} = \sum_{a=1}^{3N}\lambda_a^{-1},\end{aligned} \tag{S25}$$

where $\lambda_a$ is the a$^{th}$ eigenvalue of $\mathbf{A_1}$. Similarly, $\langle R_{ete}^2\rangle$ and the individual bond vectors can be found using the eigenvectors, $\mathbf{Q}$, from $\mathbf{A}^{-1}$:

$$\langle R_{ete}^2 \rangle = \sum_{a=1}^{3N}\left[\left(Q_{N,a}^x - Q_{1,a}^x\right)^2 + \left(Q_{N,a}^y - Q_{1,a}^y\right)^2 + \left(Q_{N,a}^z - Q_{1,a}^z\right)^2\right]\lambda_a^{-1}$$

$$\langle l_i^2 \rangle = \sum_{a=1}^{3N}\left[\left(Q_{i+1,a}^x - Q_{i,a}^x\right)^2 + \left(Q_{i+1,a}^y - Q_{i,a}^y\right)^2 + \left(Q_{i+1,a}^z - Q_{i,a}^z\right)^2\right]\lambda_a^{-1},$$

where $Q_{ia}^x, Q_{ia}^y$, and $Q_{ia}^z$ have the same meaning as in the main text, but for the eigenvectors of $\mathbf{A}_1$ instead of $\mathbf{A}$. The agreement between the structural properties calculated directly from the simulation and using the modes agree to within the numerical precision of the simulation data. Analogous expressions can be found for the modes generated with the inclusion of the hydrodynamic interaction using the eigenvectors of $\mathbf{HA}_1$, but the eigenvalues of $\mathbf{A_1}$.

### III. EXAMPLE CALCULATION FOR FINDING $\xi_{a,x}, \xi_{a,y}, \xi_{a,z}$

Here, an example calculation for finding $\xi_{a,x}(t)$, $\xi_{a,y}(t)$, and $\xi_{a,z}(t)$ for a two-bead system ($N = 2$) is illustrated. For this system, $\Delta\vec{R}(t)$ is written as

$$\Delta\vec{R}(t) = \left[\Delta x_1, \Delta y_1, \Delta z_1, \Delta x_2, \Delta y_2, \Delta z_2\right]^T.$$

Using the definition of $\xi_a(t)$, $\xi_a(t) = \sum_i Q_{ai}^{-1}\Delta\vec{R}_i(t)$, we obtain

$$\xi_a(t) = Q_{a1}^{-1}\Delta x_1(t) + Q_{a2}^{-1}\Delta y_1(t) + Q_{a3}^{-1}\Delta z_1(t) + Q_{a4}^{-1}\Delta x_2(t) + Q_{a5}^{-1}\Delta y_2(t) + Q_{a6}^{-1}\Delta z_2(t).$$

Collecting the $\Delta x_i(t), \Delta y_i(t),$ and $\Delta z_i(t)$ terms gives

$$\xi_a(t) = \left[Q_{a1}^{-1}\Delta x_1(t) + Q_{a4}^{-1}\Delta x_2(t)\right] + \left[Q_{a2}^{-1}\Delta y_1(t) + Q_{a5}^{-1}\Delta y_2(t)\right] + \left[Q_{a3}^{-1}\Delta z_1(t) + Q_{a6}^{-1}\Delta z_2(t)\right].$$

Defining

$$\xi_{a,x}(t) = Q_{a1}^{-1}\Delta x_1(t) + Q_{a4}^{-1}\Delta x_2(t) = \sum_{i=1}^{2} Q_{ai,x}^{-1}\Delta x_i(t) = \sum_{i=1}^{N} Q_{ai,x}^{-1}\Delta x_i(t),$$

$$\xi_{a,y}(t) = Q_{a2}^{-1}\Delta y_1(t) + Q_{a5}^{-1}\Delta y_2(t) = \sum_{i=1}^{2} Q_{ai,y}^{-1}\Delta y_i(t) = \sum_{i=1}^{N} Q_{ai,y}^{-1}\Delta y_i(t),$$

$$\xi_{a,z}(t) = Q_{a3}^{-1}\Delta z_1(t) + Q_{a6}^{-1}\Delta z_2(t) = \sum_{i=1}^{2} Q_{ai,z}^{-1}\Delta z_i(t) = \sum_{i=1}^{N} Q_{ai,z}^{-1}\Delta z_i(t),$$

we have

$$\xi_a(t) = \xi_{a,x}(t) + \xi_{a,y}(t) + \xi_{a,z}(t).$$

### IV. JUSTIFICATION FOR EXCLUDING $|\vec{\xi_a}|$ FROM THE LE4PD-XYZ FREE-ENERGY SURFACES

Each of the LE4PD-XYZ modes, $\vec{\xi_a}(t)$ describes a one-dimensional, collective coordinate embedded in the 3-dimensional $(x, y, z)$-Cartesian space originating at the protein's center-of-mass. In describing the dynamics of the LE4PD-XYZ modes on the two-dimensional $(\theta_a, \phi_a)$ surfaces, the radial coordinate defining the magnitude of the mode vector $|\xi_a|$ has been averaged over. That is, $F(\theta_a, \phi_a)$ can be defined using $P(\theta_a \phi_a) = \int P\left(|\vec{\xi_a}|, \theta_a, \phi_a\right) d|\vec{\xi_a}|$ as follows:

$$
\begin{aligned}
F(\theta_a, \phi_a) &= -k_B T \ln\left[P(\theta_a \phi_a)\right] \\
&= -k_B T \ln\left[\int P\left(|\vec{\xi_a}|, \theta_a, \phi_a\right) d|\vec{\xi_a}|\right]
\end{aligned}
\tag{S26}
$$

In the main text, and in previous publications involving an LE4PD analysis,[3–6] only $F(\theta_a, \phi_a)$ is reported, with the radial coordinate $|\vec{\xi_a}|$ not taken into account. Here, the use of $F(\theta_a, \phi_a)$ without explicitly including $|\vec{\xi_a}|$ in the description of the LE4PD-XYZ free-energy surfaces is justified.

Figure S1 shows the one-dimensional probability $P\left(|\vec{\xi_a}|\right)$ as a function of $|\vec{\xi_a}|$ for the first LE4PD-XYZ mode without hydrodynamics; $P\left(|\vec{\xi_a}|\right)$ has a single peak at 0.41 nm. $F(\theta_a, \phi_a)$ for three values of $|\vec{\xi_a}|$ (indicated by the vertical, dashed liens in Figure S1) are shown as subplots beneath $P\left(|\vec{\xi_a}|\right)$. In addition, the real-space fluctuations of ubiquitin described by traversing the minimum free-energy pathway between the two deepest minima on each $F(\theta_a, \phi_b)$ at a given $|\vec{\xi_a}|$ is given below the respective free-energy surface. For the smallest $|\xi_a|$ bin, the motion on the $(\theta, \phi)$ surface is mostly diffusive, with no major barriers to cross; the associated real-space fluctuations are likewise small. As $|\xi_a|$ is increased, the barriers become more well-defined until the most probable value of $|\xi_a|$ is reached. As the barriers between minima become more well-defined, the magnitude of the real-space fluctuations in the alpha-carbon sequence of ubiquitin become larger in magnitude, but describe the same type of motion as found along the $F(\theta_a, \phi_a)$ surfaces for smaller values of $|\xi_a|$. At large enough values of $|\xi_a|$, the two minima are well-localized and sampled almost exclusively, leading to the largest amplitude motions in the protein along the given mode.

This analysis demonstrates that, even though the $|\xi_a|$ is averaged over when reporting the $F(\theta_a, \phi_a)$ surfaces, the major information lost is the magnitude of the fluctuations along the given mode coordinate, as might be expected given that the coordinate averaged out is the *magnitude* of the mode vector. However, this result also indicates that the $(\theta_a, \phi_a)$ representation of the dynamics gives the correct qualitative motions of the protein along the mode coordinate, but averages out their magnitude, so the magnitude of reported motions may be slightly under- or over-estimated with respect to the 'true' fluctuations, but the discrepancy is not likely to be significant, given the mono-modal distribution of $|\xi_a|$.

### V. LE4PD-XYZ MODE DYNAMICS FROM FREE-ENERGY PATHWAYS

Shown in Figures S2, S3, and S4 are the dynamical fluctuations predicted when for LE4PD-XYZ modes 2, 5, and 7 when HI are neglected, which are three of the four slowest LE4PD-XYZ modes as predicted using the MSM analysis described in the main text. As is the case for Figure 1 in the main text, in Figures S2 and S3, the interpolated pathway shows large-scale motions in the same regions of the protein as the LE4PD-XYZ pathways (mostly the C-terminal tail region of the protein, in agreement with the LML plots of Figure 6 of the main text), but the 'details' of the motions differ considerably. For modes 2 and 5, it is also seen that, as with Figure 1 in the main text, the interpolation procedure predicts fluctuations along the entire backbone of the protein that is not seen in the LE4PD-XYZ pathway predictions or the LML for that mode. These motions seem to be an artifact induced through the use of the interpolation procedure for visualizing the mode's dynamics.

In the case for mode 7 shown in Figure S4 is slightly different, as, although the interpolated pathway does not begin or end in energetic minima nor does it follow the minimum energy pathway between minima, it still accurately predicts fluctuations in the 50 s loop of ubiquitin with some small deformations of the C-terminal tail region, in agreement with the pathways predicted by the LE4PD-XYZ method. The interpolation method for this mode also shows much less 'spurious' deformations along the backbone of ubqiuitin where no fluctuations are predicted by the corresponding LML analysis. Tentatively, since mode 7 corresponds to a large-amplitude, but highly localized fluctuation, it seems as though the interpolation approach could function well for describing this type of motion, but will generally fail at accurately describing large-amplitude, more delocalized fluctuations.

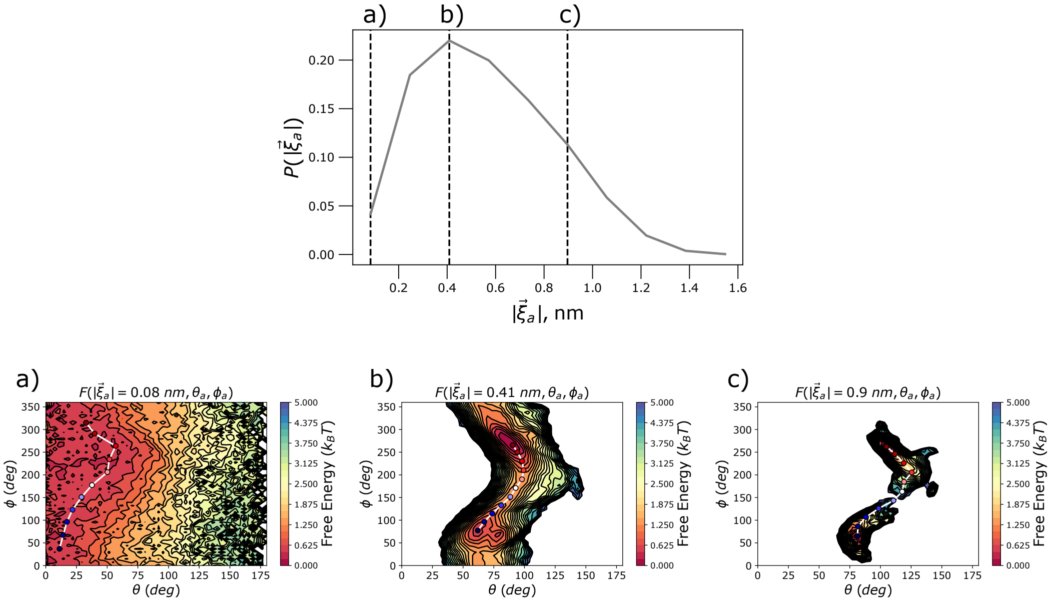


FIG. S1. Plot of the probability of sampling $|\vec{\xi}_a|$, $P\left(|\vec{\xi}_a|\right)$, as a function of $|\vec{\xi}_a|$. The $(\theta_a, \phi_a)$ free-energy surfaces for three level curves of $|\vec{\xi}_a|$ are shown for a) $|\vec{\xi}_a| = 0.08$ nm, b) $|\vec{\xi}_a| = 0.41$ nm (the most probable value of $|\vec{\xi}_a|$, and c) $|\vec{\xi}_a| = 0.9$ nm. Structural changes in ubiquitin corresponding to motion along these $(\theta_a, \phi_a)$ surfaces for each discrete value of $|\vec{\xi}_a|$ are given below the respective free-energy surfaces.

### VI. MORE ON THE OVERLAP MATRIX

Using the definition of $\xi_a(t)$ given in the Theory section, it can be seen that $O$ is the absolute value of the change of basis matrix [7] between the LE4PD-XYZ basis set without and with HI, respectively:

$$\xi_a = \sum_i Q^T_{ai} \Delta R_i = \sum_i Q^T_{ai} \sum_b Q'_{ib} \xi'_b = \sum_b \sum_i Q_{ia} Q'_{ib} \xi'_b = \sum_b \left(Q_a \cdot Q'_b\right) \xi'_b, \quad \text{(S27)}$$

where un-primed quantities are calculated without HI and primed quantities are calculated with HI. Since each eigenvector is normalized, $\max_{i,j} O_{ij} = 1$ and $O_{ij} \in [0, 1]$. This overlap metric is used to evaluate the effect of adding hydrodynamics by comparing directly the dynamics predicted by the modes in each treatment with the highest overlap between them. $O$ is calculated using only the 3N - 6 eigenvectors corresponding to the internal modes; for ubiquitin, a protein composed of 76 residues, 3N - 6 = 222 and $O$ is a 222 × 222 matrix.

Recalling that since the columns of both $\mathbf{Q}$ and $\mathbf{Q}'$ are normalized to unit magnitude, the entries of $O$ can also be

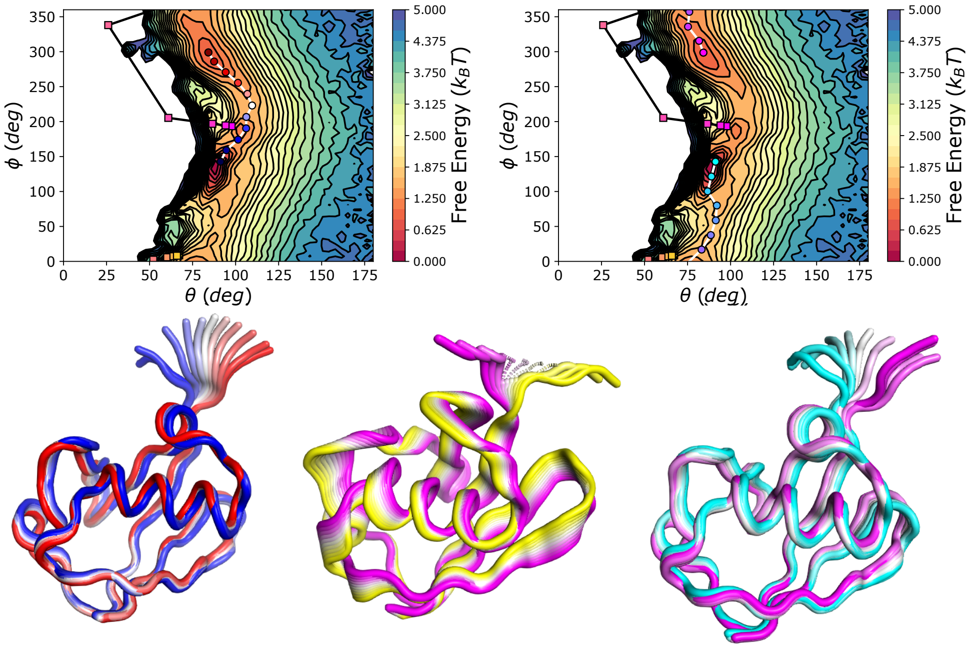


FIG. S2. Free-energy surface for the first LE4PD-XYZ mode, solved for the case where $\mathbf{H} := \mathbf{I}$, so that the LE4PD-XYZ mode solutions are identical to the calculated PCs. The surface on the left and right are identical; the blue-white-red path demonstrates one minimum energy pathway between the two minima found using the string method and the yellow-white-magenta path shows the trajectory on the surface found using the linear interpolation method between the two extreme structures along this mode coordinate. The surface on the right shows a second minimum energy pathway that crosses the periodic boundary at $\phi = 0^\circ = 360^\circ$ in cyan-white-magenta. The real-space, 3D fluctuations corresponding to each of these pathways are depicted by the ubiquitin structures given the cartoon representation below the surfaces. Each structure is colored corresponding to the analogously colored image along the pathway.

written as the cosine of the angles between the eigenvectors calculated without and with HI, $\cos(\vartheta_{ab})$, since, by the definition of the dot product,

$$Q_a \cdot Q'_b = |Q_a||Q'_b| \cos(\vartheta_{ab}) = \cos(\vartheta_{ab}) .$$

So, even for an entry in $O_{ab}$ equal to 0.9 corresponds to angle of $\vartheta_{ab} = \arccos(0.9) = 25.8^\circ$ between those two modes, which is a substantial shift in direction caused by the inclusion of hydrodynamic effects. The only modes from the two approaches that are rotated less than $10^\circ$ with respect to each other are the first two modes, whose similarity can be seen further by comparing the free-energy surfaces, sequence-localized dynamics, and timescales predicted by the first two modes in each approach.

### VII. EXAMPLE MARKOV STATE MODEL PARAMETERIZATION FOR THE LE4PD-XYZ MODES

The Markov state modeling procedure for the LE4PD-XYZ modes is nearly identical to the Markov state modeling procedure used for the isotropic LE4PD modes in [6]; all the steps to build the Markov state models (MSMs) are performed using PyEMMA. [8] For the LE4PD-XYZ MSMs, since the coordinates (or features) of the MSM have been selected *a priori*, there remain only two adjustable parameters: the MSM lagtime $\tau$ and the number of discrete states in the model $L$. Here, an example of how the MSMs analyzed in the main text are constructed is given for LE4PD-XYZ mode 7 without HI, but the same procedure is used for the other LE4PD-XYZ modes reported in the main text, regardless if HI were included or not.

To select the number of discrete states for the model, we use the VAMP-2 score [9, 10] on the first ten eigenvalues of the MSM calculated for the testing set. The VAMP is described in detail in [10], but, briefly, the VAMP divides the input data set into a training and test set, parameterizes a MSM on the training set, then evaluates its performance on the test set. The better the performance on the test set (with performance measured via the sum of the first $n$ eigenvalues of $\mathbf{T}^{test}(\tau)$ raised to some power), the higher the VAMP score and the more robust the model.

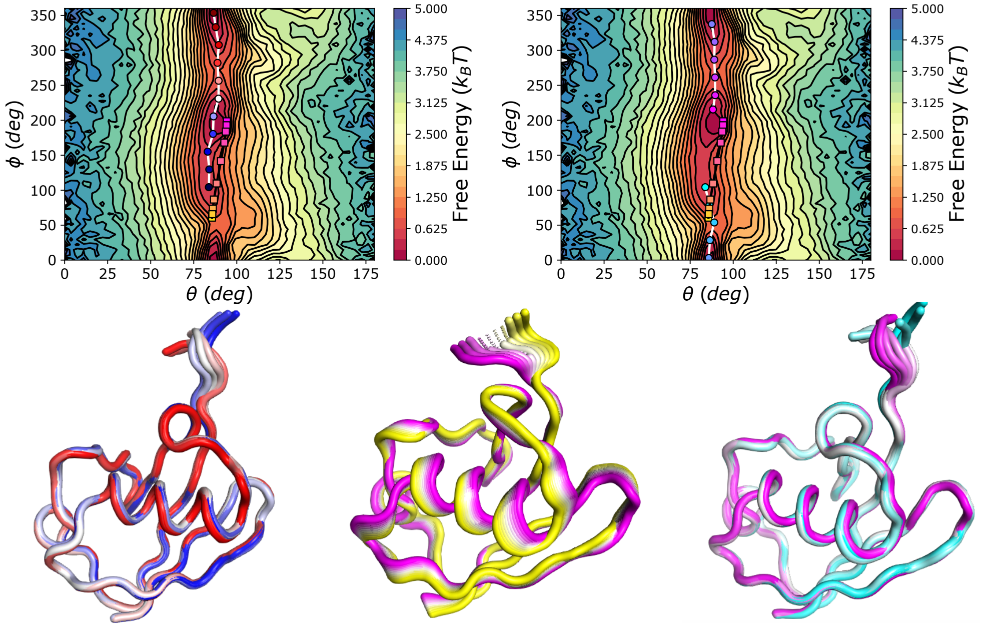


FIG. S3. Same as Figure S2, except for LE4PD-XYZ mode 5.

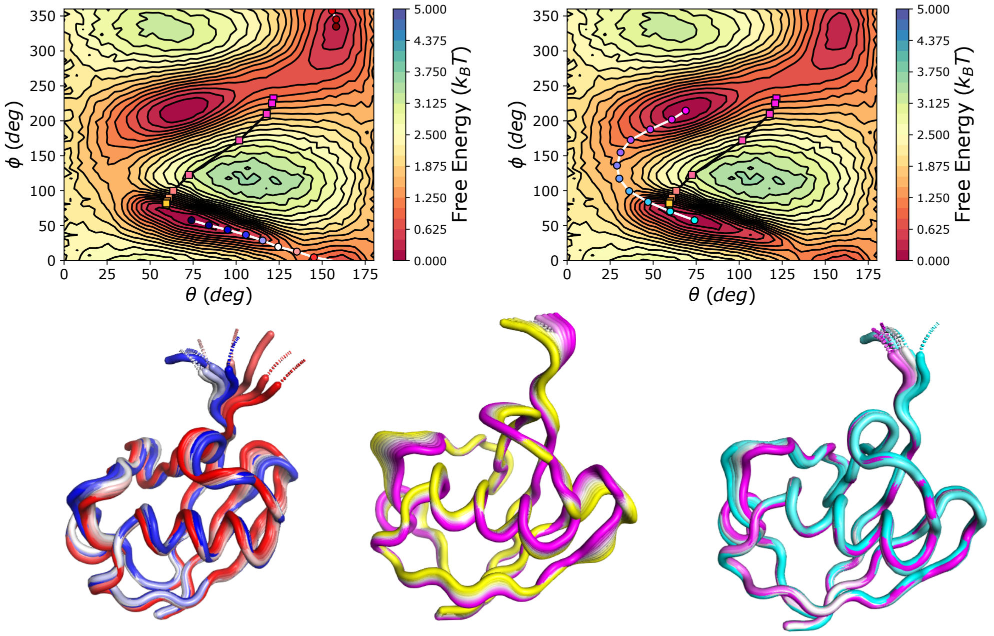


FIG. S4. Same as Figures S2 and S3, except for LE4PD-XYZ mode 7.

The VAMP-2 score for LE4PD-XYZ mode 7 without hydrodynamics is reported as the number of discrete states is varied is shown in Figure S5. The figure demonstrates that, between 100 and 5000 discrete states, the VAMP-2 score on the test set is approximately constant, within the reported uncertainty (which is the 90% confidence interval, given by the shaded region in Figure S5). Based on the data reported in Figure S5, the choice of $L = 1000$ used in the MSMs reported in the main text is justified.

For a given LE4PD-XYZ mode, the $(\theta_a, \phi_a)$ surface is discretized into $L = 1000$ states using the k-means++

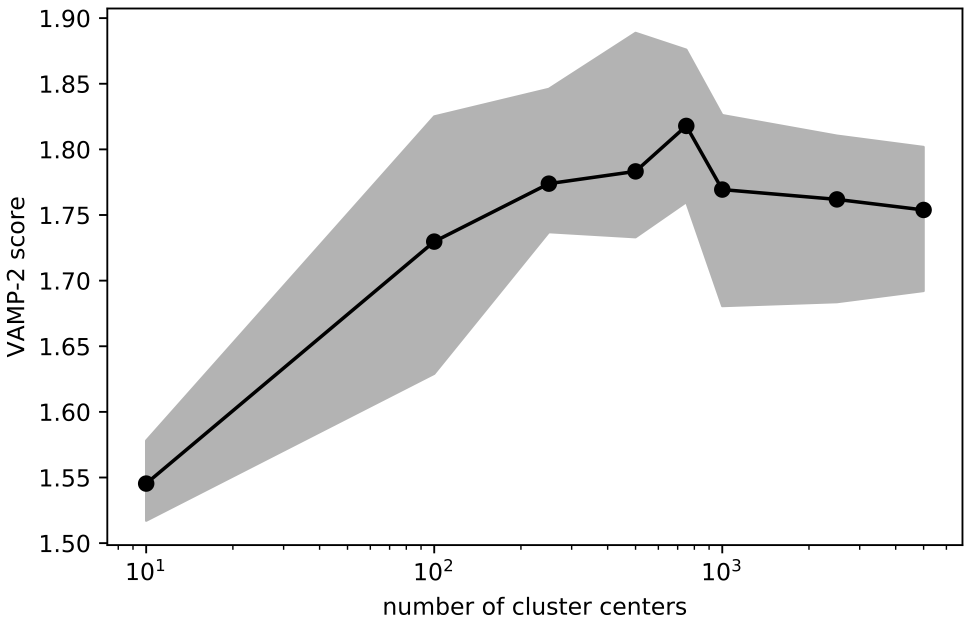


FIG. S5. VAMP-2 score as a function of $L$ (labled as 'number of cluster centers' on the abscissa of the plot) for LE4PD-XYZ mode. Shaded areas give the 90% confidence interval.

algorithm. [11] The lagtime of each MSM is determined using the procedure described in the main text, where the longest $\tau$ such that the maximum and minimum projection of $\psi_2$ from the model remain in the deepest minima on the $(\theta_a, \phi_a)$ surface. The transition matrix, $\mathbf{T}(\tau)$, is constructed using the reversible, maximum likelihood estimator given in [12]:

$$T_{ij} = \frac{(c_{ij} + c_{ji})\,\pi_j}{c_i\pi_j + c_j\pi_i}, \tag{S28}$$

with $c_{ij}$ the number of observed transitions from discrete state $i$ to discrete state $j$, $c_i = \sum_j c_{ij}$ the total number of observed transitions from state $i$ to all other states $j$ after a lagtime $\tau$ has elapsed, and $\pi_i$ is the stationary probability of state $i$, i.e. what fraction of the total simulation time does the system reside in state $i$. This transition matrix is diagonalized to obtain the eigenvalues $\lambda_i^{\mathrm{MSM}(\tau)}$ and left and right eigenvectors $\phi_i$ and $\psi_i$, respectively. Using the relationship between the relationship between $\mathbf{T}(\tau)$ and its corresponding rate matrix, $\mathbf{K}(\tau)$[13], $\mathbf{T}(\tau) = e^{\mathbf{K}(\tau)\tau}$, the timescales of the $i^{th}$ process of the MSM can be determined as [14, 15]

$$t_i = -\frac{\tau}{\ln\left(\lambda_i^{\mathrm{MSM}(\tau)}\right)}.$$

Since $\lambda_1^{\mathrm{MSM}(\tau)} = 1$, $t_1 = \infty$, which corresponds to the 'trivial' long-time, equilibrium state. So, the slowest process of the MSM is given by $t_2$; a plot of $t_2$ as a function of the MSM lagtime $\tau$ is shown in Figure S6 for LE4PD-XYZ mode 7, without HI.

Here, we are only interested in the slowest timescale from the MSM on the $(\theta_a, \phi_a)$ surfaces, which is described by $\psi_2$. Figure S7 shows the projection of $\psi_2$ onto the $(\theta_a, \phi_a)$ surface for LE4PD-XYZ mode 7 without HI; in this case, the process describes the flow of probability density from the top two minima on the surface to the minimum around $(\theta_a, \phi_a) \approx (75, 60)$ $(deg)$.

[1] L. Landau and E. Lifshitz, *Fluid Mechanics: Volume 6* (Elsevier Science, 2013).
[2] E. Caballero-Manrique, J. K. Bray, W. A. Deutschman, F. W. Dahlquist, and M. G. Guenza, Biophysical Journal **93**, 4128 (2007).

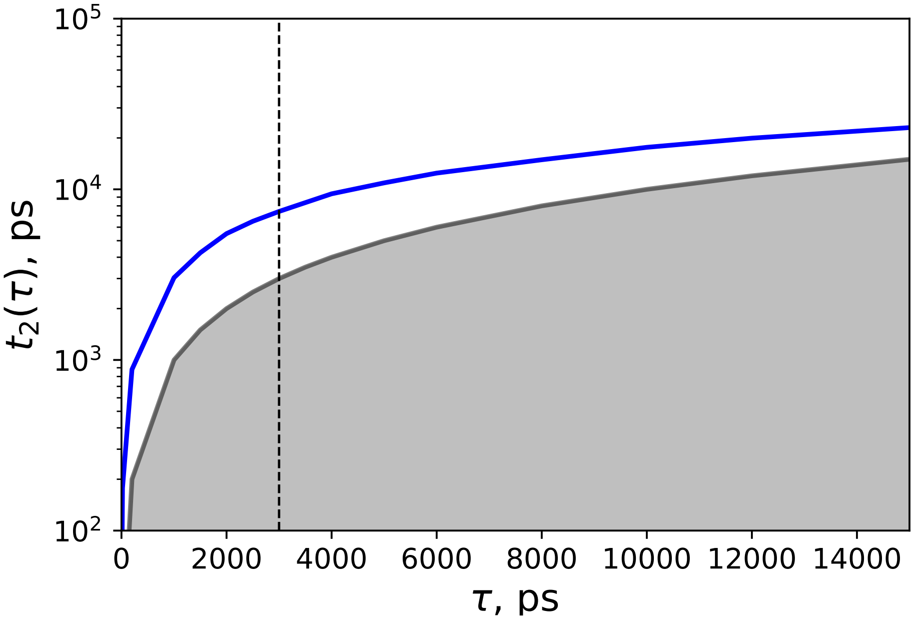


FIG. S6. Plot of $t_2$ as a function of $\tau$, the MSM lagtime, for LE4PD-XYZ mode 7 without HI. The vertical, dashed line shows the lagtime at which the MSM analyzed for this study was constructed. The grey, shaded area shows the region where $t_2 \leq \tau$, where the given process relaxes faster than the observation interval, and the process cannot be resolved by the MSM.

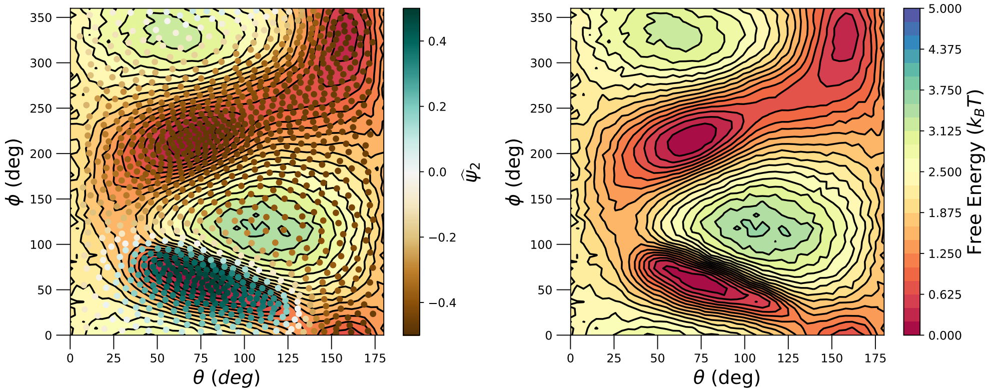


FIG. S7. Left: Plot of a scaled version of $\psi_2$, $\widehat{\psi}_2 = \frac{\psi_2 - \min(\psi_2)}{\max(\psi_2) - \min(\psi_2)} - 0.5$, projected onto the $(\theta, \phi)$ surface for LE4PD-XYZ mode 7 without HI. Right: Free-energy surface alone for LE4PD-XYZ mode 7 without HI.

[3] J. Copperman and M. G. Guenza, Journal of Physical Chemistry B **119**, 9195 (2015).
[4] J. Copperman and M. G. Guenza, Journal of Chemical Physics **145**, 015101 (2016).
[5] J. Copperman, M. Dinpajooh, E. R. Beyerle, and M. G. Guenza, Physical Review Letters **119**, 158101 (2017).
[6] E. R. Beyerle and M. G. Guenza, The Journal of Chemical Physics **151**, 164119 (2019), https://doi.org/10.1063/1.5123513.
[7] R. A. Horn and C. R. Johnson, *Matrix Analysis* (Cambridge University Press, New York, NY, USA, 1986).
[8] M. K. Scherer, B. Trendelkamp-Schroer, F. Paul, G. Pérez-Hernández, M. Hoffmann, N. Plattner, C. Wehmeyer, J. H. Prinz, and F. Noé, Journal of Chemical Theory and Computation **11**, 5525 (2015).
[9] H. Wu and F. Noé, arXiv preprint arXiv:1707.04659 (2017).
[10] M. K. Scherer, B. E. Husic, M. Hoffmann, F. Paul, H. Wu, and F. Noé, The Journal of Chemical Physics **150**, 194108 (2019).
[11] D. Arthur and S. Vassilvitskii, Proceedings of the eighteenth annual ACM-SIAM symposium on discrete algorithms **8**, 1027 (2007).
[12] B. Trendelkamp-Schroer, H. Wu, F. Paul, and F. Noé, Journal of Chemical Physics **143**, 174101 (2015).

[13] L. Reichl, *A Modern Course in Statistical Physics* (Wiley, 1998).
[14] W. C. Swope, J. W. Pitera, and F. Suits, The Journal of Physical Chemistry B **108**, 6571 (2004).
[15] G. Bowman, V. Pande, and F. Noé, *An Introduction to Markov State Models and Their Application to Long Timescale Molecular Simulation*, Advances in Experimental Medicine and Biology (Springer Netherlands, 2013).
